# The Canadian Pediatric Imaging Platform (C-PIP) Reference Sample: Neuroimaging Protocol and Cross-site Validation for Harmonized Multimodal Pediatric Imaging

**DOI:** 10.64898/2026.10.26.761379

**Authors:** Mervyn Singh, Shireen Parimoo, Vicente Enguix, Filomeno Cortese, Milton Camacho, Sean Spinney, Samantha J. Lynch, Helen L. Carlson, Lara M. Leijser, Christine L. Tardif, Ilana R. Leppert, G. Bruce Pike, Emile Kadalie, Jamie Near, Chathura Kumaragamage, Riya Chitroda, Michel Louis Lauzon, Kathryn Y. Manning, Gopal Varma, Tomas Paus, Ashley D. Harris, Gregory A. Lodygensky, Miriam H. Beauchamp, Donald J. Mabbott, Daniel C. Kopala-Sibley, Jacob A. S. Vorstman, Keith Owen Yeates, Paul D. Arnold, Sébastien Jacquemont, Darren S. Kadis, Sarah Lippé, Lune Bellec, Catherine Lebel, Patricia Conrod, Signe Bray, Anne L. Wheeler

## Abstract

Understanding how the human brain matures from infancy through adolescence is essential for identifying factors that influence health outcomes. Magnetic resonance imaging (MRI) provides powerful tools for characterizing structural, microstructural, functional, and metabolic aspects of brain development; however, many pediatric imaging studies are constrained by small sample sizes, limited geographic diversity, lack of longitudinal measurement, restricted age ranges, and heterogeneous acquisition protocols. While recent large-scale neuroimaging consortia have begun to address these challenges, such resources remain limited in the Canadian context, creating a critical gap. The Canadian Pediatric Imaging Platform (C-PIP) has developed a multisite neuroimaging framework across three of Canada’s largest pediatric hospitals: Alberta Children’s Hospital (Calgary), Centre Hospitalier Universitaire Sainte-Justine (Montréal), and the Hospital for Sick Children (Toronto). These three sites were united through a national initiative, leveraging high-performance MRI systems to accelerate Canadian neurodevelopmental research. In addition, C-PIP has implemented a centralized data sharing infrastructure to facilitate open science and collaborative research across institutions. Here, we describe the development and implementation of a multimodal MRI acquisition protocol designed to support multimodal longitudinal studies of typical and atypical brain development spanning infancy through adolescence. Within-site stability and cross-site stability of C-PIP core MRI sequences were evaluated using a travelling-participant study. The C-PIP MRI protocol establishes a harmonized, multisite imaging framework that supports large-scale investigations of pediatric brain development and disorders.

**Highlights:**

1. The Canadian Pediatric Imaging Platform (C-PIP) is a national initiative acquiring longitudinal phenotyping and neuroimaging data from 820 children aged 0-18 years across three Canadian provinces (Alberta, Ontario and Québec).
2. C-PIP has developed a harmonized, developmentally sensitive neuroimaging protocol to collect high-quality, comprehensive imaging data across multiple modalities.
3. This paper outlines the development, considerations and implementation of the C-PIP neuroimaging protocol across sites.
4. Preliminary cross-site validation analyses demonstrate good within- and between-site stability in neuroimaging measures.

**Graphical Abstract:** 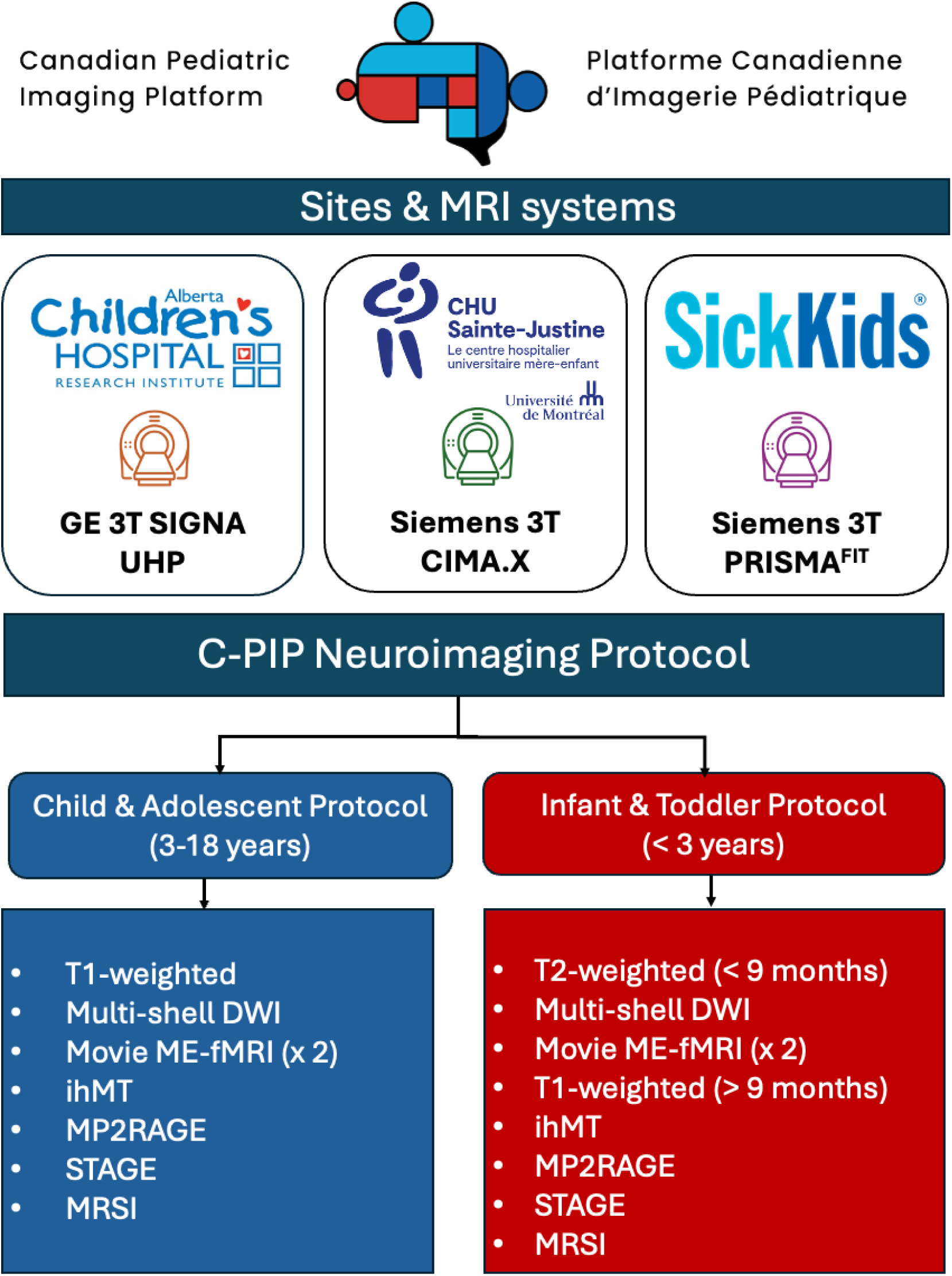

## 1. Introduction

Disruptions to normative brain development during childhood are important risk factors for a variety of conditions throughout the lifespan, from intellectual disabilities and psychiatric disorders emerging in early childhood and adolescence (Kretzer et al., 2024; Parkes et al., 2021; Paus et al., 2008; Reisinger and Hannan, 2025), to neurodegeneration and dementia in later life (Schor and Bianchi, 2021; Shabani and Hassan, 2023). Longitudinal magnetic resonance imaging (MRI) can be used to track brain development and identify critical windows where disruptions might influence developmental outcomes (Bethlehem et al., 2022).

Large-scale multisite neuroimaging initiatives have begun to address these important questions by coordinating acquisition protocols and pooling data across research centers. Programs such as the Adolescent Brain Cognitive Development (ABCD) Study (Casey et al., 2018), the Human Connectome Project–Development (HCP-D) (Somerville et al., 2018), and the HEALthy Brain and Child Development (HBCD) Study (Dean et al., 2024; Volkow et al., 2024) demonstrate the value of harmonized imaging protocols and shared infrastructure for studying brain development. However, several important gaps remain. Although large Canadian youth-specific datasets exist, such as the Saguenay Youth Study (Pausova et al., 2017) and the Province of Ontario Neurodevelopmental Disorders Network (POND) (https://pond-network.ca/), longitudinal data spanning the full spectrum of pediatric development from infancy through adolescence are limited. Further, current large-scale developmental datasets are predominantly U.S.-centric, thus restricting geographic, social, and public health policy representativeness.

The Canadian Pediatric Imaging Platform (C-PIP) unites three of Canada’s largest pediatric hospital research centres: Alberta Children’s Hospital (ACH; Calgary), Centre Hospitalier Universitaire Sainte-Justine (CHUSJ; Montréal), and the Hospital for Sick Children (SickKids - SK; Toronto). C-PIP provides a unique opportunity to characterize neurodevelopment across a wide age range, aiming to recruit a reference cohort of 820 participants across the three sites, spanning infancy through adolescence, with annual follow up visits for up to two years post baseline.

In this paper, we (1) describe the development and implementation of a harmonized C-PIP MRI protocol used in older children and adolescents and a modified protocol for children younger than 3 years of age across three pediatric imaging centers, and (2) report initial cross-site stability analyses based on a travelling-participant study designed to quantify within-site and between-site repeatability of core imaging measures. Other publications will describe the concept and design of the C-PIP platform [Bray et al., under review], reference sample recruitment and phenotyping [Lynch et al., under review], and bioinformatics infrastructure [Spinney et al., under review] in more detail.

## 2. The C-PIP MRI Protocol

### 2.1 Protocol Design and Considerations

The C-PIP MRI protocol was developed to characterize normative patterns of brain development in a Canadian-specific reference cohort spanning infancy to late adolescence. The protocol was developed by an MRI working group involving investigators, MRI physicists, and imaging scientists from participating sites. An extensive testing and validation process was implemented through an iterative procedure of sequence development, cross-site harmonization on key contrast-related parameters, initial piloting of scans, and preliminary qualitative and quantitative comparisons. The guiding principles for protocol development balance scientific rigor with feasibility in pediatric populations while enabling harmonized data acquisition across sites with different scanners and vendors.

To enable comprehensive high-quality measurements of brain structure and function, a multimodal protocol was emphasized that included T1- and T2-weighted structural imaging (prioritized according to the age of the child, see Section 3.3.1 for details), multi-shell diffusion MRI, and multi-echo functional MRI. These sequences were designated as core acquisitions due to their broad uptake within the neuroscientific community (Gozdas et al., 2018; Lebel et al., 2019; Mills et al., 2016), and, given the pervasiveness of head motion in pediatric populations (Afacan et al., 2016; Thomson et al., 2024), were prioritized early in the scanning session when children are at their most compliant to maximize high-quality data. Innovative techniques, including myelin-sensitive imaging (inhomogeneous magnetization transfer imaging; ihMT), quantitative T1 mapping (magnetization prepared 2-rapid gradient echo imaging; MP2RAGE), multiparametric quantitative imaging (strategically acquired gradient echo imaging; STAGE), and whole-slice magnetic resonance spectroscopic imaging (MRSI) were included as additional “extended” acquisitions to provide complementary measures of brain tissue microstructure, composition, and metabolism (Chen et al., 2018; Marques et al., 2010; Varma et al., 2015; Zhu and Barker, 2011). The inclusion of these latter acquisitions was designed to support ongoing methodological innovation and enable the validation of novel neuroimaging biomarkers within a harmonized multisite framework.

Given the wide age-range of investigation in C-PIP (0-18 years), the MRI protocol was designed to be sensitive to developmentally varying aspects of tissue contrast and physiology (Spann et al., 2023; Turesky et al., 2021). As head motion is higher in pediatric populations, sequences were selected to be as short (≤10 min) and efficient as possible, minimizing the impact of motion-related artifacts on data quality while retaining data integrity. To increase feasibility across the entire developmental range represented in C-PIP, two age-appropriate versions of the protocol were created, consisting of a “Child and Adolescent” protocol (scan age: 3-18 years) and an “Infant and Toddler” protocol (scan age: < 3 years), with the latter characterized by shorter sequences given the higher propensity for motion in younger children (Afacan et al., 2016; Pardoe et al., 2016; Thomson et al., 2024) and optimized parameters (see Section 3, Table 2).

**Table 1.**
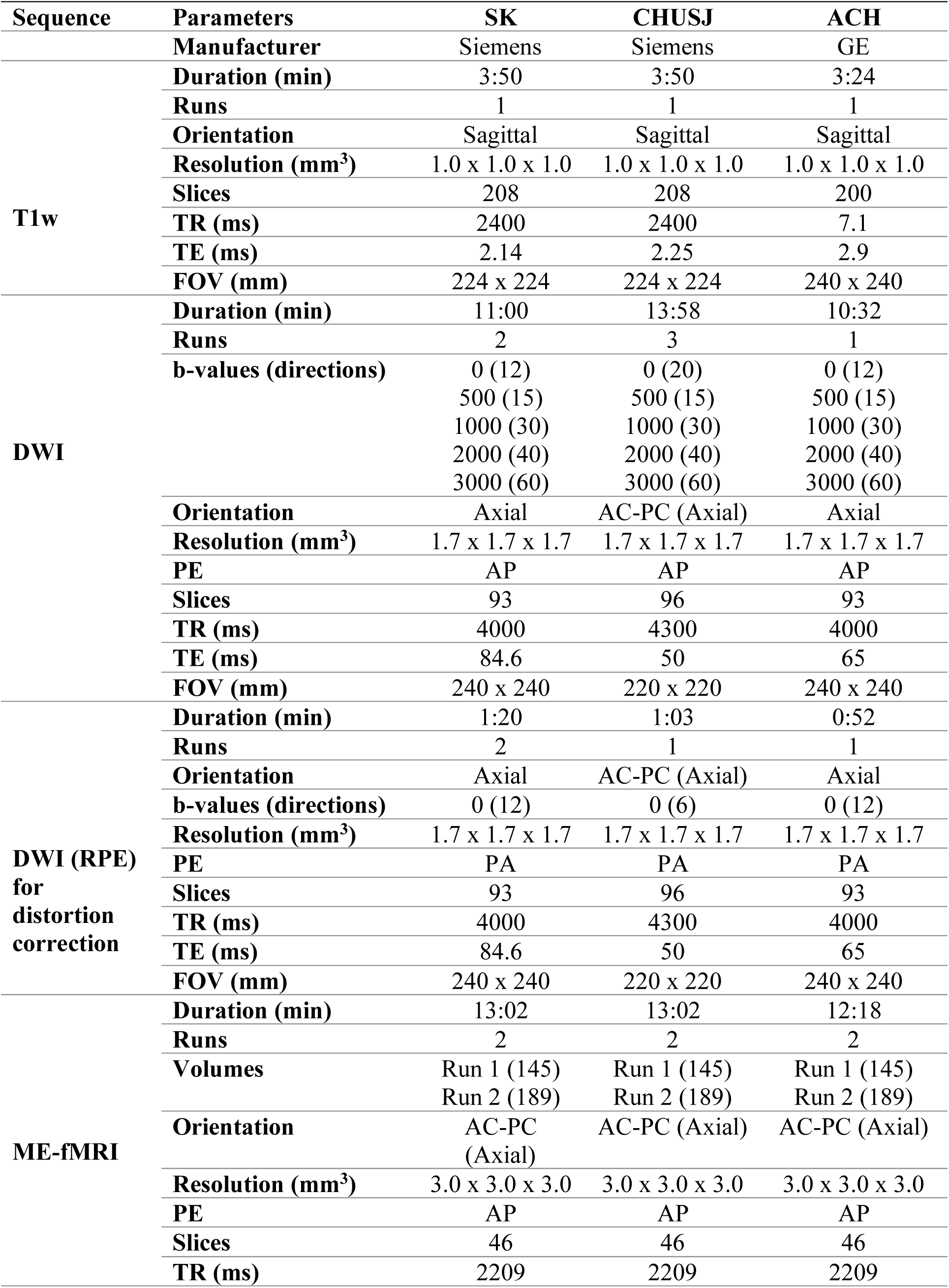
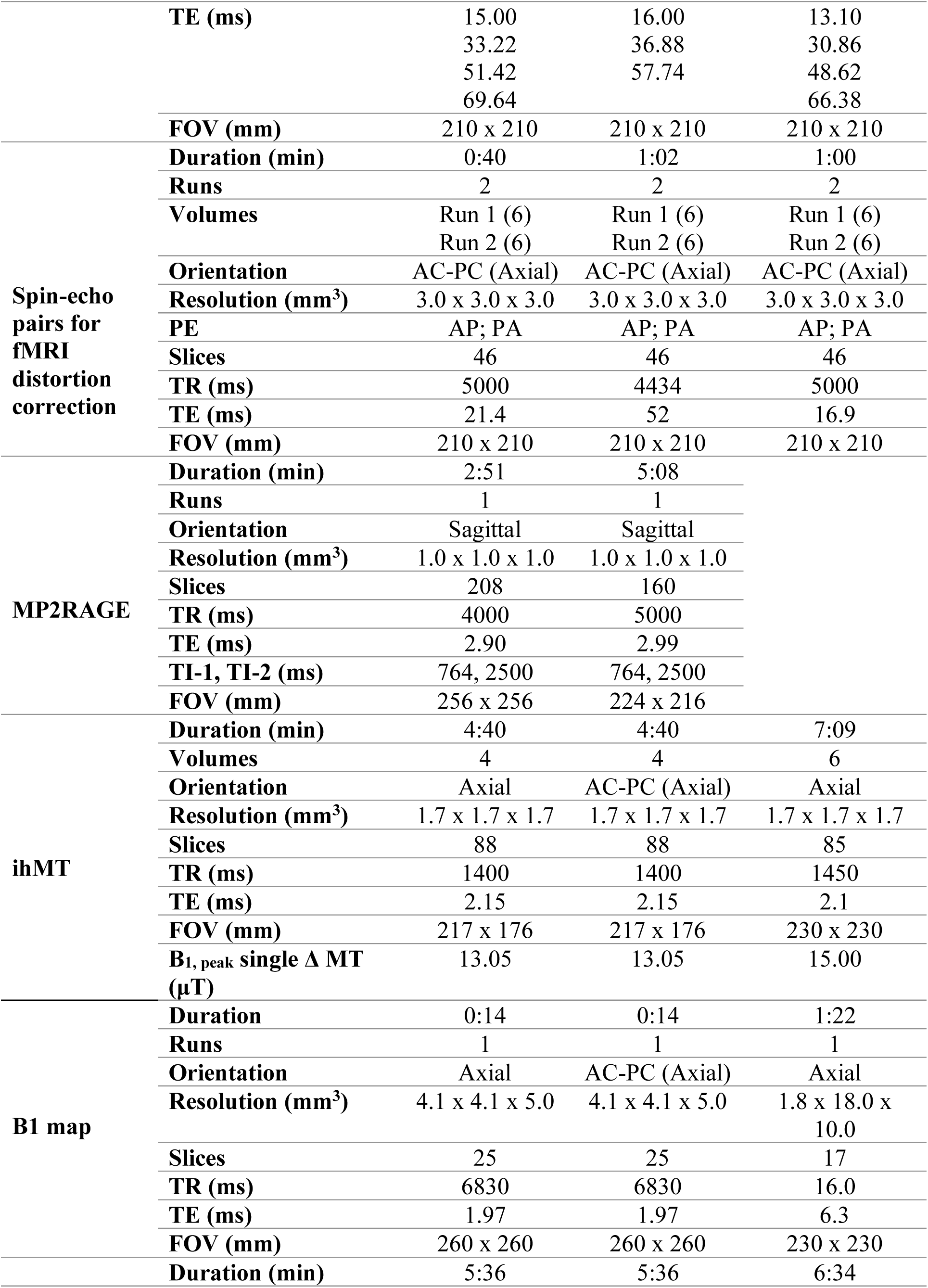
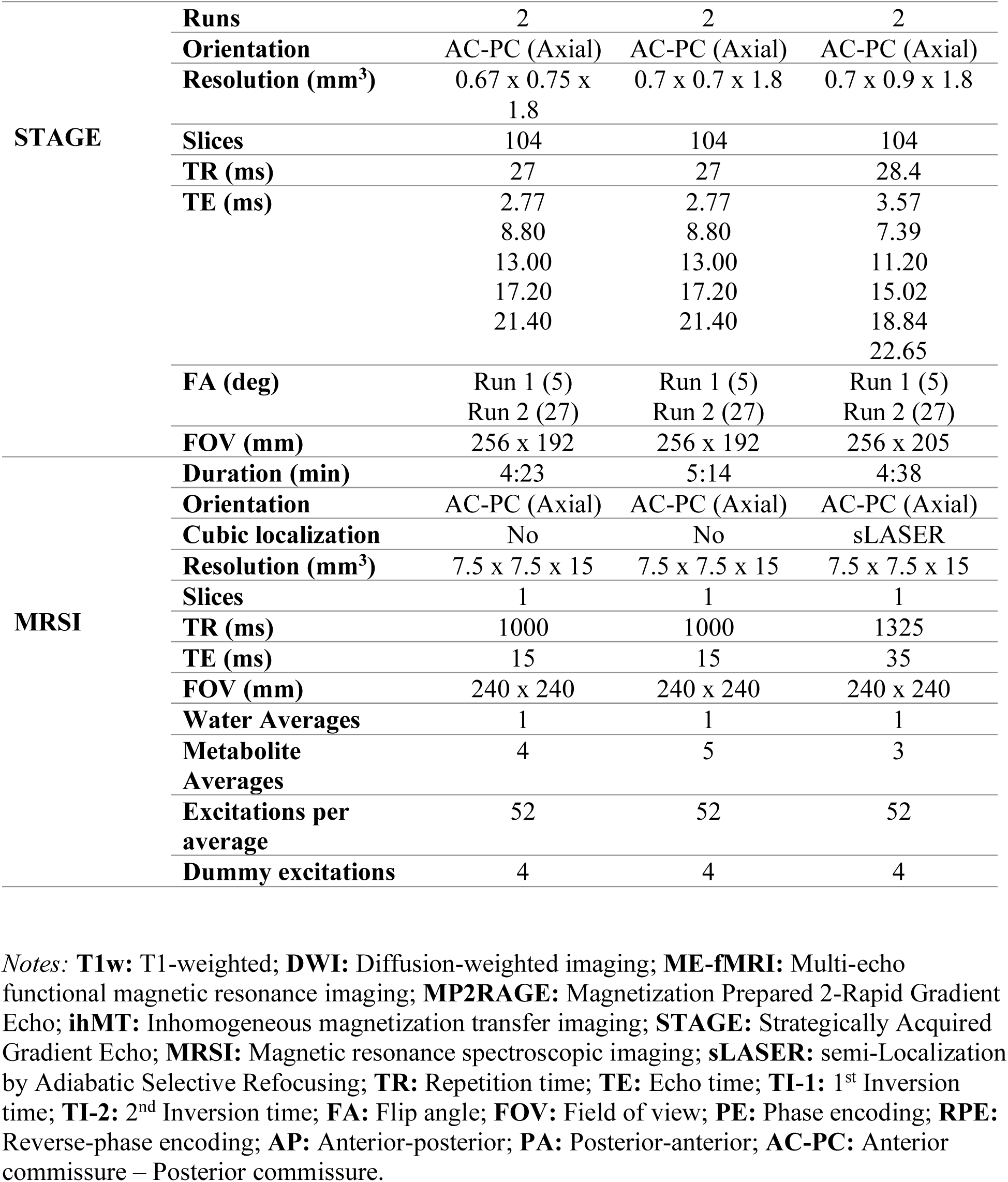
C-PIP Child and Adolescent Protocol.

**Table 2.**
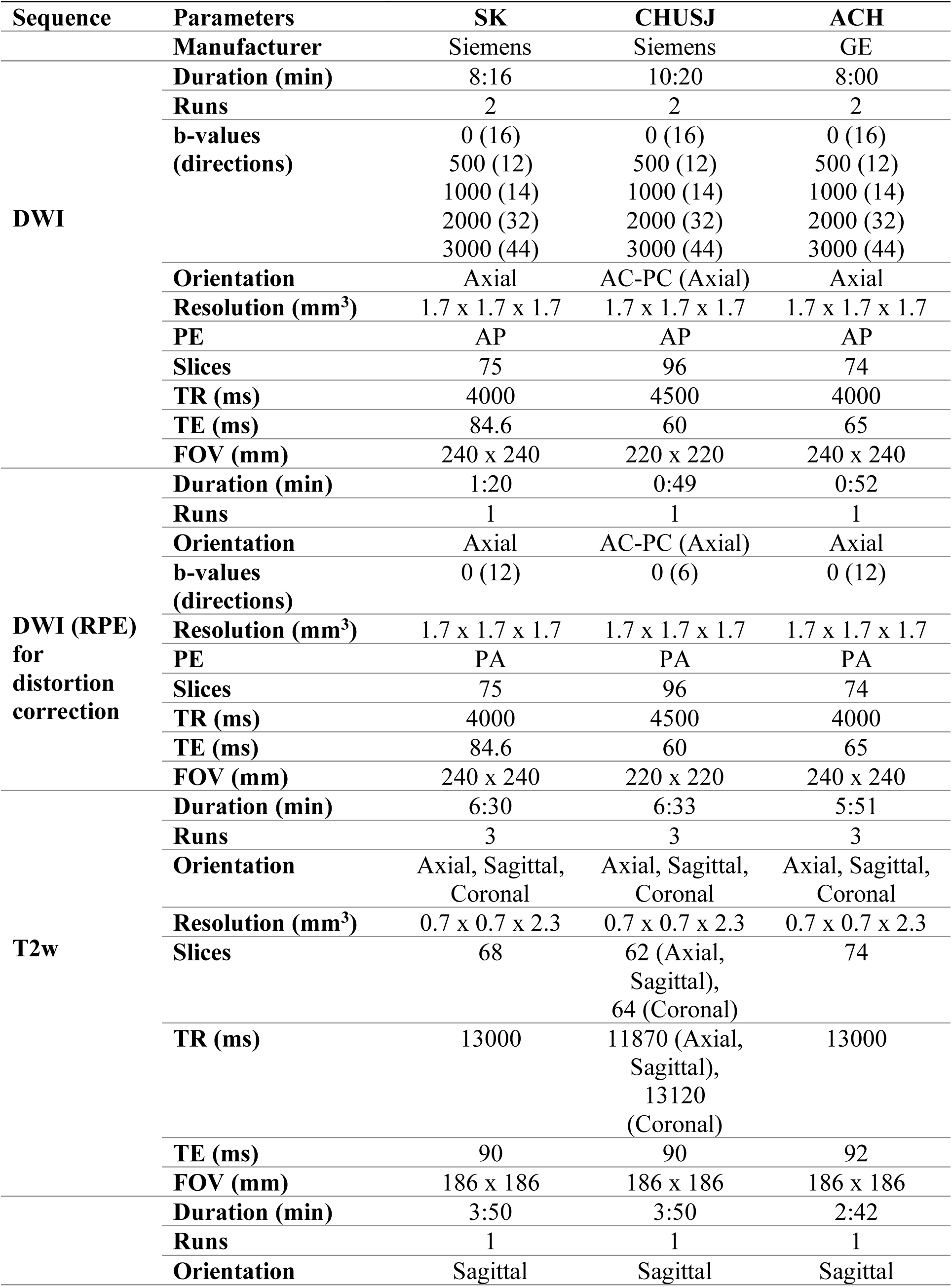
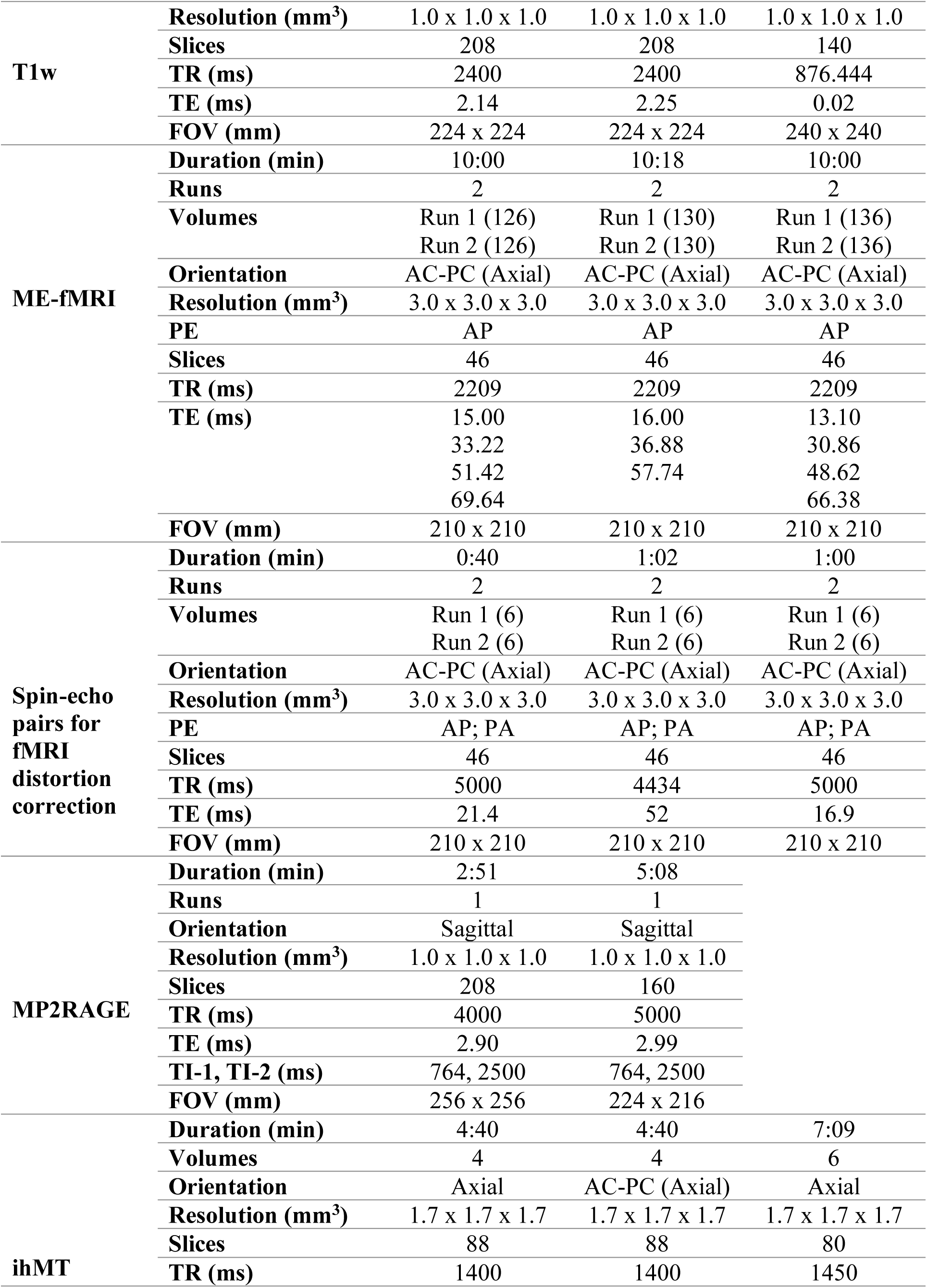
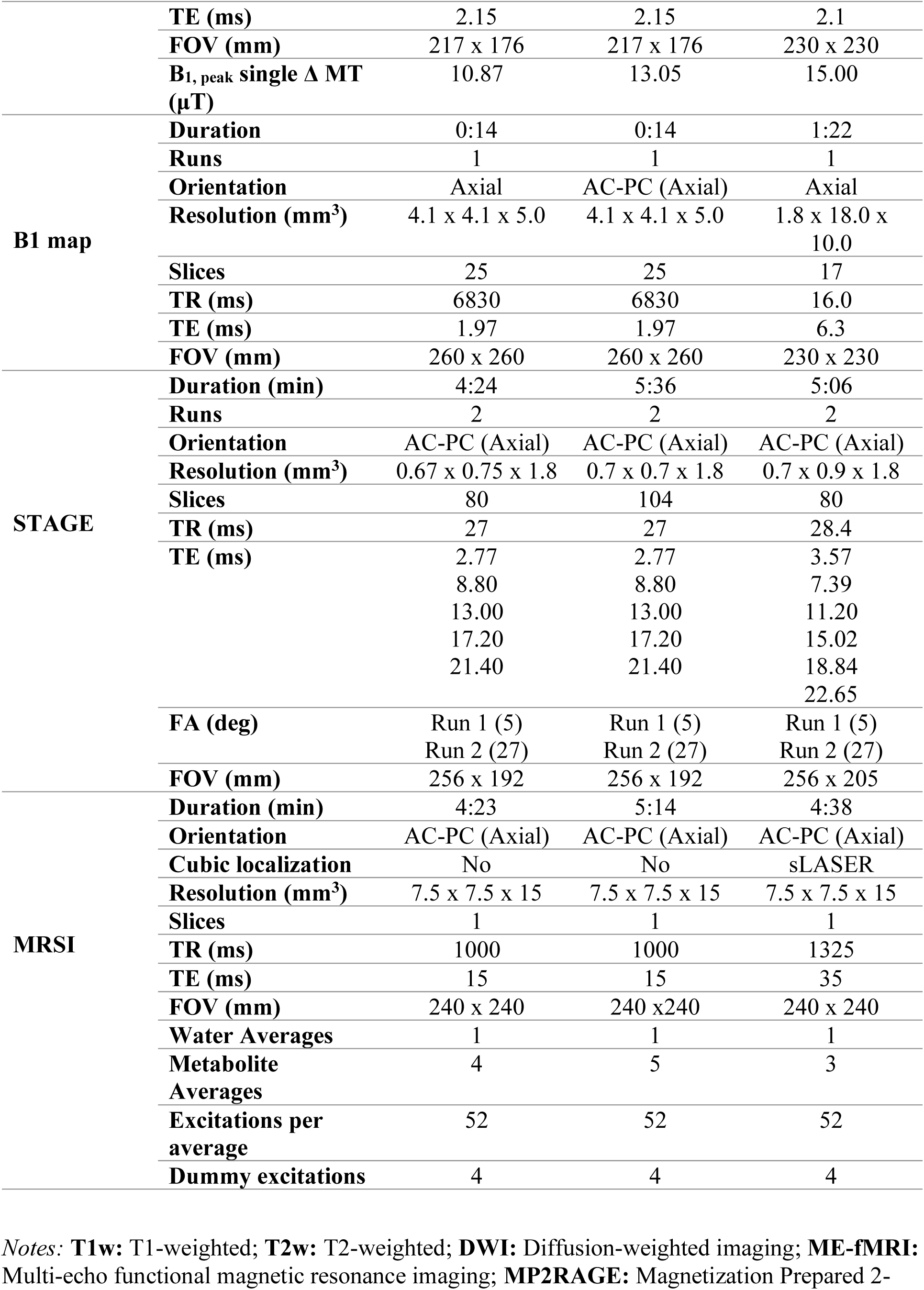
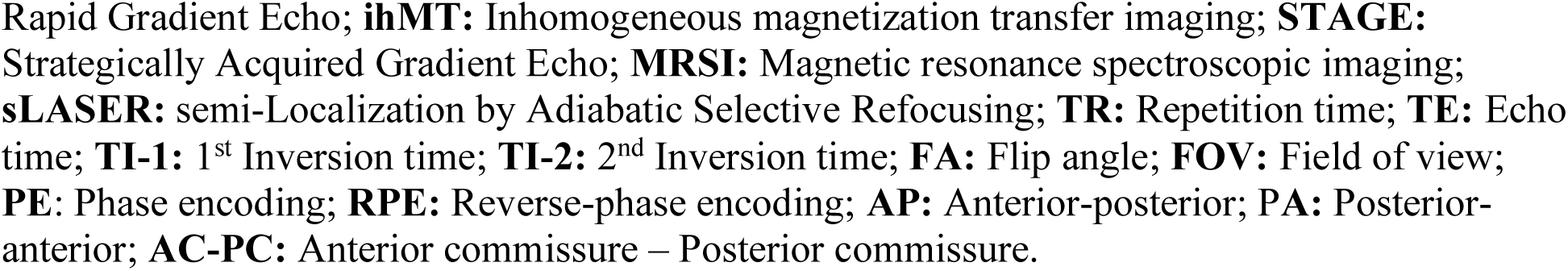
C-PIP Infant and Toddler protocol.

Achieving comparability across sites is complex and requires careful consideration of cross-vendor differences in hardware capabilities. While a minimum of technical alignment is required, a too stringent approach will decrease feasibility of collaboration. Hence, rather than strictly enforcing identical acquisition parameters across scanner platforms, protocol development prioritized alignment of key imaging characteristics, such as spatial resolution, diffusion weighting, and acquisition geometry, while allowing vendor-specific optimizations such as echo time (TE) and repetition time (TR) for some sequences. For example, while the TR for the diffusion-weighted imaging sequence in the Child and Adolescent protocol differed between sites, all sites were matched on spatial resolution (1.7 x 1.7 x 1.7 mm^3^), number of diffusion directions (145), and b-values (*b* = 500, 1000, 2000, 3000 s/mm^2^). Such an approach allows each scanner platform to operate within its technical capabilities while maintaining comparable data quality and minimizing unnecessary increases in scan duration. Collectively, this balanced approach facilitates robust multicenter data integration while accommodating the unavoidable differences in scanner hardware and software across imaging centers.

Lastly, we should note that, while the C-PIP MRI protocol was designed to remain stable throughout data acquisition, minor adjustments to sequence parameters were made towards the start of data collection for optimization purposes. These changes have been documented. Over time, further minor adjustments to sequence parameters may occur as scanners are upgraded or new acquisition techniques become available. Any further modifications will be documented and evaluated to ensure compatibility with previously acquired datasets. Together, the C-PIP protocol provides a harmonized imaging strategy spanning early infancy through adolescence, supporting longitudinal investigation of brain development.

## 3 Protocol Details

This section provides a detailed description of the individual MRI sequences included in the C-PIP protocol, outlining their acquisition parameters, decisions and optimizations made during protocol development, and role within the overall imaging framework. Representative images from each modality are presented in Supplementary Figures 1-8. Details of the major sequence parameters for each protocol across sites are provided in Tables 1 (Child and Adolescent) and 2 (Infant and Toddler). A Flowchart of the sequences included for both protocols is presented in Figure 1.

**Figure 1.**
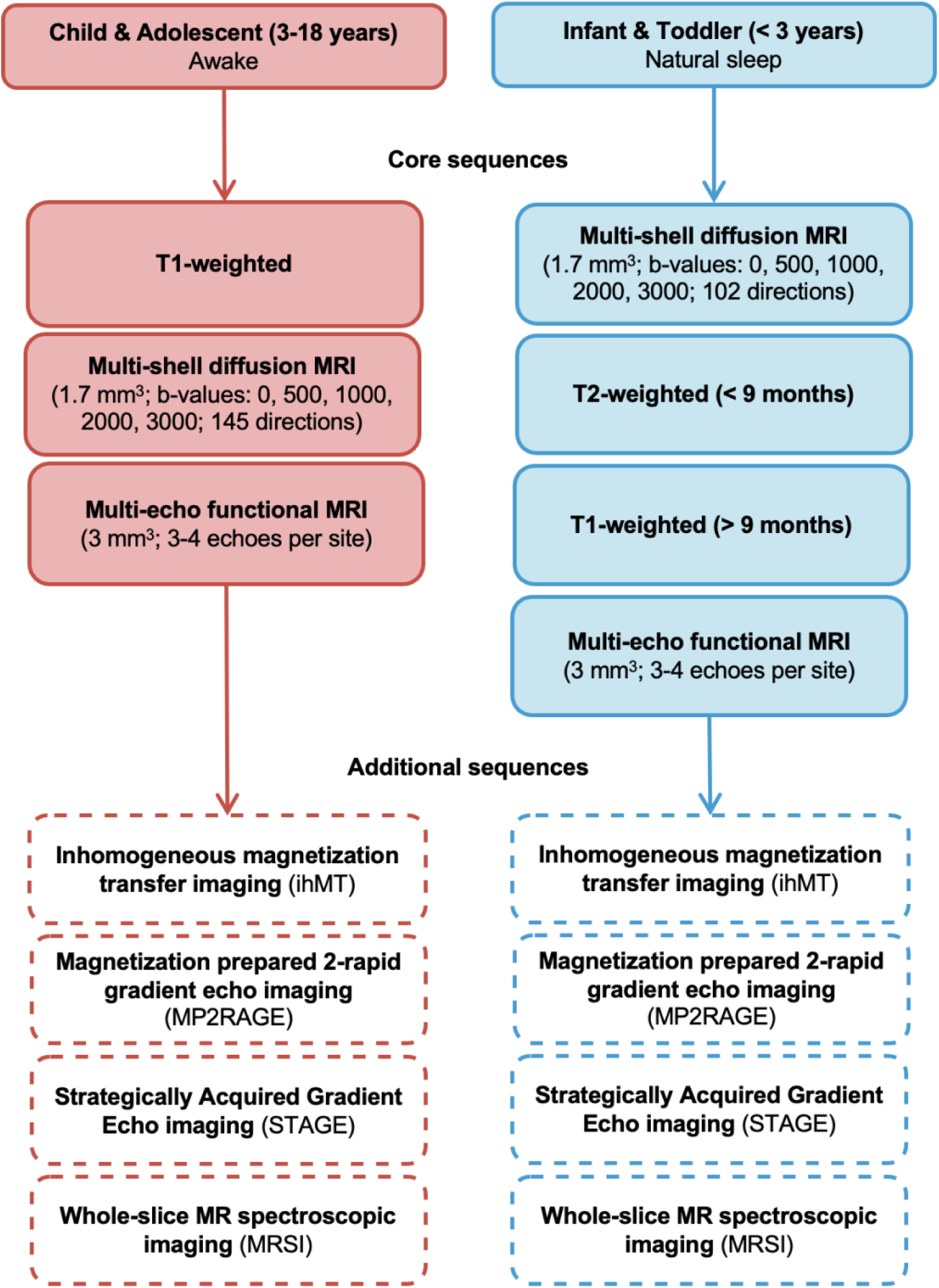
Flowchart depicting the neuroimaging sequences included in the C-PIP Child and Adolescent and Infant and Toddler protocols.

### 3.1 Scan Procedure

Given the challenges of obtaining high-quality data in pediatric populations, several strategies are employed across sites to maximize participant comfort and engagement during scanning. Prior to the actual scan session, children aged 3 years and older are provided with child-friendly resources including two YouTube videos, *“MRI Brain Scan”* (Madisonhealth, 2021) and *“MRI Sounds Inside Scan Room”* (MRI & CT Scan Tech, 2023), and a picture book, “*Pluto and the MRI Rocket Ship Adventure*” (Frayne, 2015), that contain information about MRI and what participants can expect from a typical scanning session. While these videos are presented in English, there is the option for audio dubbing in the participants preferred language (e.g., French) in cases where English is not their first language. In addition, both ACH and CHUSJ have a mock MRI scanner that mimics the experience of an actual scanning session. Participants at these sites are given the opportunity to practice in the mock scanner prior to the scanning session to acclimate to the scanning environment. During the scan, participants are permitted to watch an age-appropriate movie or television show, usually chosen from a preselected list of G and PG movies, or through a streaming service (i.e., Disney Plus, Netflix, YouTube).. For functional MRI sequences, a standardized movie stimuli specified in the study protocol is used (Section 3.3.3). To minimize variability related to first-time exposure to the movie stimuli (Andric et al., 2016), participants are required to watch the movies before scanning.

For participants younger than 3 years old, scans are scheduled to align as closely as possible with natural sleep times. Care is taken to minimize disturbances throughout the session such as dimming the lights within the MRI room and playing white noise through the intercom to create a conducive environment for sleep. Where possible, research staff are also encouraged to book a separate room to provide parents the opportunity to feed, swaddle and settle infants before the scanning session (King et al., 2023). When natural sleep is not possible, alternative strategies are employed to keep children engaged during scanning, including playing a video of their choice or having a parent in the MRI room to comfort them. Specialized equipment for infant scanning, such as mouldable earplugs, MiniMuffs® Neonatal Noise Attenuators (Pacific Biomedical, Scottsdale, AZ, USA), and swaddling blankets are used to provide additional protection and comfort.

The total scan duration for each protocol was targeted to last no longer than approximately 60 minutes. However, booking windows were adjusted based on the participant’s age and whether the scan was performed during the day or evening. For children 3 years and older undergoing awake scans, a 60-minute MRI booking was generally considered sufficient. For toddlers undergoing awake scans, a longer booking window of approximately 75-90 minutes was recommended to allow additional time for preparation and settling. These flexible booking windows allowed sufficient time to accommodate age-related differences, participant preparation, and individual variability in scanning readiness.

Detailed notes of all age-appropriate MRI familiarization strategies are documented for each session. In addition to providing important information on the success of these strategies for each participant and age group, these notes also assist the study team in planning for follow-up visits. In the long term, these notes may serve as a reference for other research groups to maximize their success of attaining high-quality pediatric neuroimaging data.

### 3.2 MRI scanning systems

MRI data are collected at three pediatric hospitals using research-dedicated 3 Tesla (3T) MRI systems: a General Electric (GE) SIGNA Ultra High-Performance system (ACH – Calgary; software version: MR30.1), a Siemens MAGNETOM Cima.X (CHUSJ – Montréal; software version: XA61), and a Siemens Prisma^Fit^ (SK – Toronto; software version: XA30). Scans are acquired using the standard head-coil configuration for each scanner: a 48-channel head coil at ACH and 32-channel head coils at CHUSJ and SK. Standard safety procedures, including participant screening for MRI contraindications and use of MRI-compatible safety equipment (i.e., earplugs), are implemented across all scanning sites. Standard procedures for incidental findings were followed for each site. At SK, all baseline structural scans (i.e., T1- and/or T2-weighted) are reviewed by an experienced radiologist, and subsequent follow-up scans are sent for review only if the first scan was flagged. At ACH and CHUSJ, only structural scans flagged by clinical technologists or study staff are reviewed by a radiologist.

### 3.3 Core Sequences

#### 3.3.1 Structural MRI

Structural imaging allows for morphometric analyses (Dima et al., 2022; Frangou et al., 2022; Mills et al., 2016) and multimodal registration, thus representing a foundational aspect for any neuroimaging protocol. The structural protocol for C-PIP included a 3D T1-weighted (T1w) volume acquired sagittally at an isotropic spatial resolution of 1 mm (Supplementary Figure 1). Higher spatial resolutions (0.9 mm or 0.8 mm isotropic) were considered during protocol development; however, 1 mm resolution was selected to balance image quality (signal-to-noise ratio, SNR) with reduced scan time and more robust segmentation with tools such as FreeSurfer (Fischl, 2012). T1w scans on the Siemens platforms were acquired using a Magnetization Prepared Rapid Gradient Echo (MPRAGE) sequence, while the GE platform used the Brain Volume Imaging (BRAVO) sequence. Additionally, to minimize acoustic noise when scanning toddlers during natural sleep, a SILENZ™ version of the T1w sequence was used for the GE site.

For the Infant and Toddler protocol, T2-weighted (T2w) imaging was used (in addition to T1w) due to differences in gray-white matter contrast in newborns/young infants compared with older children (Turesky et al., 2021) (Supplementary Figure 2). At this early stage of life, the water-to-lipid ratio is inverted due to incomplete myelination (Zhang et al., 2019). Consequently, standard T1w sequences yield poor gray-white matter contrast in newborns, making tissue segmentation more challenging (Dubois et al., 2021). While both T1w and T2w sequences are included in the Infant and Toddler protocol, T2w is prioritized for children aged 9 months or younger, whereas T1w imaging is preferred for older participants. To maximize data quality, a novel acquisition strategy was used in which three short 2D scans (∼2 minutes each at 0.7 x 0.7 x 2.3 mm^3^) were acquired separately, with each scan oriented along one of the three orthogonal planes (axial, sagittal, coronal). During post-processing, a custom model-based super-resolution reconstruction algorithm is employed to combine each individual scan to generate a super-resolved 3D image with sub-millimeter resolution (0.7 mm isotropic; Askin Incebacak et al., 2022). This approach provides greater flexibility in the presence of head motion by avoiding reacquisition of the entire sequence. If motion affects only one scan from a specific orientation plane, only that scan is repeated, thereby reducing unnecessary scan time while maximizing data retention.

#### 3.3.2 Diffusion weighted Imaging (DWI)

Diffusion-weighted imaging (DWI) exploits the orientational dependency of water molecules within highly myelinated white matter fibers to characterize brain structural connectivity (Baliyan et al., 2016; Beaulieu, 2014; Rowe et al., 2016). Due to the sensitivity of DWI to various aspects of tissue microstructure, this approach has been widely used to investigate white matter maturational profiles in typical (Genc et al., 2018; Grotheer et al., 2022; Lebel et al., 2012; Reynolds et al., 2019; Schilling et al., 2023; Singh et al., 2026) and atypical (Chen et al., 2016; Dimond et al., 2019; Kelly et al., 2016; Kirkovski et al., 2024) neurodevelopment.

To provide detailed, biologically specific characterization of white-matter microstructure in pediatric populations (Dhollander et al., 2021; Pines et al., 2020; Tournier et al., 2013; Tuch et al., 2002; Zhang et al., 2012), the C-PIP MRI protocol implemented a high-angular resolution multi-shell sequence (Supplementary Figure 3). Diffusion data for C-PIP was collected in the axial plane at an isotropic spatial resolution of 1.7 mm. Images were collected orthogonally at ACH and SK, whereas at CHUSJ, images were aligned along the anterior and posterior commissural planes (AC-PC). The acquisition includes 145 diffusion-encoding directions in the Child and Adolescent protocol and 102 directions in the Infant and Toddler protocol spread across 4 b-values (*b* = 500, 1000, 2000, and 3000 s/mm²). In addition to the main diffusion acquisition, 12 to 20 *b* = 0 s/mm^2^ images are also collected, interspersed among the other images. A separate series of 6 to 12 reverse-phase encoded *b* = 0 s/mm² images was collected in the posterior-anterior (P-A) direction to correct for susceptibility distortions during preprocessing (Andersson et al., 2003). Two reverse-phase encoded series were collected at SK (one for each run), while at ACH and CHUSJ, only one pair was acquired.

This protocol was adapted from the diffusion sequence implemented in the ABCD study, with modifications to optimize diffusion modelling in younger populations. Specifically, the C-PIP sequence incorporates denser angular sampling at lower b-values (see below) to improve the stability of diffusion tensor and multi-compartment microstructural estimates (Assaf and Basser, 2005; Genc et al., 2020; Jeurissen et al., 2014; Jones, 2004; Zhang et al., 2012). Whereas the ABCD protocol acquires 6 directions at *b =* 500 s/mm², 15 directions at *b* =1000 s/mm², and 15 directions at *b* = 2000 s/mm², the C-PIP Child and Adolescent protocol acquires 15 directions at *b* = 500 s/mm², 30 directions at *b* = 1000 s/mm², and 40 directions at *b* = 2000 s/mm². The diffusion sequence was further optimized in the Infant and Toddler protocol to ensure scan feasibility while maintaining data quality (12 directions at *b =* 500 s/mm², 14 directions at *b =* 1000 s/mm², 32 directions at *b* = 2000 s/mm², and 44 directions at *b* = 3000 s/mm²).

Consideration was given whether to acquire all b-values in a single run or split across multiple runs. Acquiring all b-values in a single run minimizes variability in between-run motion but also increases the risk of incomplete data if interrupted mid-scan. In contrast, splitting the acquisition across multiple runs preserves data from completed runs if later acquisitions fail, although it introduces additional scan time and between-run variability. The split-run approach also provides a practical advantage by allowing the acquisition to be divided into shorter, more manageable segments, reducing the impact of within-run motion and enabling earlier completion of lower b-value acquisitions. For the Child and Adolescent protocol, the diffusion sequence was split across separate runs at the Siemens sites. At CHUSJ, three runs were acquired: Run 1 (*b* = 500 s/mm², 15 directions; *b* = 1000 s/mm², 30 directions), Run 2 (*b* = 2000 s/mm², 40 directions), and Run 3 (*b* = 3000 s/mm², 60 directions). At SK, the acquisition was split across two runs, each including *b* = 500 s/mm² (Run 1: 8 directions, Run 2: 7 directions), *b* = 1000 s/mm² (Run 1: 14 directions, Run 2: 16 directions), *b* = 2000 s/mm² (Run 1: 21 directions, Run 2: 19 directions), and *b* = 3000 s/mm² (Run 1: 29 directions, Run 2: 31 directions). The sequence implemented at the ACH/GE site was collected in a single run, with all b-values interspersed throughout the acquisition. For the Infant and Toddler protocol, images were acquired across two runs (51 directions per run) for all sites, which enables data from a completed first run to be retained if an infant wakes or moves before the second run.

Importantly, the GE system employs a “TR-cycling” approach, whereby rather than acquiring all slices for an entire volume sequentially before moving to the next volume, the acquisition collects a given slice for every volume in a cycle before proceeding to the next slice. This approach enables shorter TE and TR but invalidates typical motion correction strategies. We limited volume cycling to 2 TRs, in which slices are acquired in an interleaved fashion across pairs of volumes. This retains some of the sequence optimization of full-volume TR-cycling (i.e., short TE and TR) but allows for coarse motion correction to still be applied during post-processing, with the exception of slice-to-volume motion correction via the “eddy” function in FSL (Andersson et al., 2017; Bastiani et al., 2019).

#### 3.3.3 Multi-echo Functional MRI (ME-fMRI)

***Sequence details and parameters:*** Functional MRI (fMRI) assesses changes in the Blood Oxygenation Level Dependent (BOLD) response as a proxy of neural activity between brain regions (Buxton, 2013; Glover, 2011; Ogawa, 2012). In the C-PIP MRI protocol, fMRI is acquired across two separate runs using a multiband, multi-echo, echo-planar imaging sequence (ME-fMRI) (Supplementary Figure 4). ME-fMRI acquires images at multiple echo times (TE) (Kundu et al., 2017, 2012) and provides several advantages over conventional single-echo acquisitions, including enhanced denoising possibilities, signal recovery, and increased stability of derived measures (Constable et al., 2026; Kundu et al., 2017; Lynch et al., 2020; Moser et al., 2025; Rai et al., 2025). In the absence of a consensus on the optimal number of echoes, a maximum of 4 echoes was chosen for each fMRI run to balance scan quality and acquisition time. Vendor-specific considerations at CHUSJ dictated a 3-echo acquisition.

ME-fMRI runs were acquired axially at an isotropic spatial resolution of 3 mm, with an anterior– posterior (AP) phase-encoding direction oriented to align with the anterior and posterior commissures (AC-PC). To correct susceptibility-induced distortions, an additional pair of spin-echo images are acquired using identical spatial parameters and reversed phase-encoding directions (AP/PA) (Morgan et al., 2004). At the Siemens sites (CHUSJ and SK), ME-fMRI acquisitions also include 10 dummy volumes collected before the movie play is triggered by the scanner to allow magnetization to reach a steady state. The GE UHP system currently does not support dummy volume acquisition for this sequence, and so to achieve a steady-state magnetization, a corresponding set of volumes must be discarded manually during processing. Note that to retain alignment between temporal features across sites, volumes would need to be removed consistently across sites (despite this only being necessary at the GE/ACH site).

Functional MRI acquisitions in the Infant and Toddler protocol were shortened (2 x 5 minutes) relative to the Child and Adolescent protocol (approximately 13 minutes across 2 runs) to reduce scan duration and improve feasibility.

***Experimental paradigm and stimulus delivery:*** Naturalistic movie paradigms were chosen to improve engagement and reduce head motion, relative to traditional resting-state acquisitions (Finn and Bandettini, 2021; Vanderwal et al., 2019). In the Child and Adolescent protocol, participants view two short, Pixar animated films, Partly Cloudy (Sohn, 2009) and La Luna (Casarosa, 2012) during scanning for a total duration of approximately 13 minutes. This duration was selected to balance data quality with the need to limit overall scan time. While age-varying or participant-selected content was considered to further enhance compliance, a standardized set of films was ultimately selected to ensure consistency across participants and sites. The selected films provide engaging, high-quality socio-emotional content that is appropriate across the broad pediatric age range. Partly Cloudy has been used in prior pediatric neuroimaging studies (Kamps et al., 2022; Richardson et al., 2018), providing the opportunity to replicate findings across cohorts. Importantly, both films contain no spoken language, an important consideration for maintaining consistency across English- and French-speaking participants. Participants viewed the movies prior to scanning to reduce variability associated with first-time exposure to the movie stimulus (Andric et al., 2016) across longitudinal visits. Movie paradigms were not used for fMRI runs in the Infant and Toddler protocol, as scans are acquired during natural sleep.

However, if a child is awake during scanning, the scan will proceed with a video of their choice played to help maintain their attention. Acquisition state is recorded in the scan notes for each session. These notes also capture if a participant in the Child and Adolescent protocol fell asleep during the functional series.

At ACH, the movies were played through an MRI-compatible BOLDscreen 32-inch UHD system monitor (Cambridge Research Systems, Rochester, UK) connected to a stimulus PC located in the MRI console room. Audio was delivered through an Optoactive III audio system with a FOMRI microphone (Optoacoustics Ltd, Mazor, Israel). Movie playback was implemented using a custom-built script in PsychoPy (version 2024.1.5) (Peirce et al., 2019), which interfaced with a Lumina LSC-400B stimulus response controller (Cedrus, San Pedro, CA, USA) to automatically trigger the movie stimuli at the onset of the first volume. Movies were delivered at SK using the same MRI-compatible monitor (BOLDscreen 32-inch UHD system, Cambridge Research Systems, Rochester, UK) connected to an external laptop, with audio delivered through external speakers in the scanning room. Movies were presented using PsychoPy (version 2024.1.4) with manual triggering by the experimenter after dummy volume collection. For younger participants, SK employs an alternative audiovisual system (VisuaStim; Resonance Technology Inc. Northridge, CA, USA) that includes headphones and goggles fitted specifically for smaller head circumferences. Lastly, for CHUSJ, movies were presented through the NordicNeuroLab “fMRI Essentials Package” system (Bergen, Norway), which included an MRI-compatible 40-inch LCD monitor connected to a stimulus presentation PC, with audio playback delivered through over-the-ear-headphones. Movie playback was automatically triggered after dummy volume collection via a stimulus synchronization device (“SyncBox”) using the nordicAktiva stimulus presentation software (version 2.0.1).

### 3.4 Additional Sequences

#### 3.4.1 Inhomogeneous magnetization transfer imaging (ihMT)

Inhomogeneous magnetization transfer (ihMT) imaging was included in the C-PIP protocol to provide a myelin-sensitive contrast that complements the structural, diffusion, and relaxometry acquisitions (Alsop et al., 2023; Geeraert et al., 2018; Munsch et al., 2021; Varma et al., 2015) (Supplementary Figure 5). Magnetization transfer (MT) imaging generates myelin-sensitive contrast by exploiting interactions between free water and macromolecular proton pools within the tissue. IhMT differs from conventional MT imaging by selectively detecting signals arising from long-lived dipolar order within macromolecular structures (Manning et al., 2017; Swanson et al., 2017; Varma et al., 2017). Because these dipolar-order effects are prominent in the highly ordered lipid membranes of myelin, ihMT exhibits greater specificity to myelin than conventional MT imaging (Alsop et al., 2023; Duhamel et al., 2019). During protocol development, MT saturation (MTsat) was considered as an alternative myelin-sensitive imaging technique (Helms et al., 2008). Although MTsat offers higher SNR, higher resolution, and shorter scan times, ihMT was selected because current evidence suggests it provides greater biological specificity for myelin, making it well suited for studying neurodevelopment where progressive myelination is a major feature of brain maturation (de Faria et al., 2021; Deoni et al., 2012; Dipnall et al., 2024; Geeraert et al., 2018; Grotheer et al., 2022).

A custom acquisition based on a 3D ihMTRAGE sequence described by Varma et al., (2020) was implemented on Siemens and GE scanners to support cross-site consistency. This includes 1 s MT preparations of 5-ms, Tukey-shaped MT pulses (frequency offset Δ = 7 kHz) every 100 ms before 88 gradient echo readouts (flip angles: 8°) of a segmented radial fan beam acquisition.

The type of MT preparation acquired for ihMT (zero power, +Δ, −Δ, or dual Δ achieved by cosine modulation) is changed after acquisition of a complete volume. At the Siemens sites (CHUSJ and SK), a total of four volumes were collected: (1) dual Δ, (2) +Δ, (3) −Δ, and (4) zero-power reference. At the GE site (ACH), an additional repetition of the dual Δ image was collected to improve SNR. In addition, to enable correction for transmit-field inhomogeneity and longitudinal relaxation effects in downstream quantitative analyses, the protocol includes the acquisition of a B1 map at all sites (Chung et al., 2010; Sacolick et al., 2010), a T1 map using MP2RAGE at the Siemens sites, and specifically for the GE site, a high flip angle on-resonance preparation (Munsch et al., 2021), thereby improving the accuracy and comparability of ihMT-derived measures across scanner platforms and individuals.

Although a lower spatial resolution (e.g., 2 mm isotropic) would have improved signal-to-noise ratio and reduced scan time, a resolution of 1.7 mm isotropic was ultimately selected to minimize Gibbs ringing and to match the diffusion MRI acquisition, facilitating voxel-wise multimodal analyses such as MRI-based *g*-ratio mapping (Berg et al., 2022; Lu et al., 2025; Mohammadi and Callaghan, 2021; Stikov et al., 2015). Sequence parameters were adjusted separately for the Infant and Toddler protocol to reduce participant burden and ensure compliance with specific absorption rate (SAR) restrictions for infants during scanning (Table 2). Specifically, at ACH, the number of slices was reduced to shorten scan time from 7:35 min to 7:09 min. Furthermore, due to SAR-related issues in young participants (0-3 years old) at SK, the radiofrequency peak amplitude (B_1, peak_) of MT preparations were decreased, with B_1,peak_ of MT pulses for single Δ lowered from 13 to 11 μT (Table 2).

#### 3.4.2 Quantitative imaging

In comparison to metrics derived from conventional MRI techniques, which are based on signal intensity differences across tissue types, quantitative MRI (qMRI) methods estimate tissue parameters in absolute physical units, thus providing biomarkers that are comparable across individuals (Radunsky et al., 2024; Saltarelli et al., 2025; Weiskopf et al., 2021). Despite not being commonly used in standard pediatric neuroimaging, qMRI had shown steady adoption for investigation of brain maturation in healthy and clinical samples (Gräfe et al., 2021; Plank et al., 2025; Schulze et al., 2025; Xu et al., 2025). To leverage the advantages offered by qMRI, the C-PIP MRI protocol includes two specific acquisition approaches.

As a complementary technique to standard T1-weighted structural imaging, a 3D Magnetization Prepared 2-Rapid Gradient Echo (MP2RAGE) volume (Supplementary Figure 6) was acquired sagittally to generate T1 relaxation maps that are sensitive to different aspects of tissue microstructure, such as myelination and axon diameter (Eminian et al., 2018; Ho et al., 2024; Kühne et al., 2021; Marques et al., 2010; Plank et al., 2025). Although MP2RAGE was considered as a potential replacement for T1-weighted imaging, concerns regarding motion sensitivity and the robustness of downstream segmentation led to its inclusion as an additional rather than a core structural acquisition. At SK, the MP2RAGE sequence was acquired using compressed sensing to reduce scan time. Implementation of MP2RAGE also posed technical challenges on the GE platform at ACH and was therefore included only at the SK and CHUSJ sites.

C-PIP also includes Strategically Acquired Gradient Echo (STAGE) imaging (Chen et al., 2018; Haacke et al., 2020) (Supplementary Figure 7), an innovative multi-echo acquisition that generates multiple quantitative MRI outputs, including T1 maps, proton density (PD) (Zrinzo et al., 2026), R2* (Raab et al., 2022), and quantitative susceptibility maps (QSM) (Liu et al., 2015; Pacchiano et al., 2026). STAGE provides an efficient and flexible framework for deriving multiple quantitative biomarkers in a short amount of time (∼6 mins total), providing comprehensive information on tissue composition, and supports advanced multimodal analyses (Saltarelli et al., 2025). STAGE data at each site was acquired across two runs, with differing flip angles (Run 1: 5°, Run 2: 27°). At the GE site, a total of 6 echoes was collected per run, while 5 echoes were acquired at the Siemens sites. Slices were acquired axially in 2D with AC-PC alignment. Post-processing and reconstruction of quantitative maps generated from STAGE rely on proprietary software tools obtained through an ongoing partnership with SpinTech MRI (Haacke et al., 2024; https://spintechmri.com/). Due to inter-subject variability in tissue relaxation properties, especially at different ages (Deoni, 2010), a separate B1 map is acquired (see section 3.4.1) and incorporated into downstream processing to enable subject-specific correction and improve quantitative accuracy.

#### 3.4.3 Magnetic resonance spectroscopic imaging (MRSI)

Magnetic resonance spectroscopic imaging (MRSI) was included to measure metabolic concentrations within the brain (Supplementary Figure 8). MRSI expands upon single-voxel MR spectroscopy by increasing spatial coverage to map the distributions of metabolite levels across a slice, thereby offering enhanced sensitivity and specificity for assessing metabolic status across different tissue types (Maudsley et al., 2020; Posse et al., 2013; Wilson et al., 2019; Zhu and Barker, 2011).

At the Siemens sites (CHUSJ and SK), a 2D single-slice Spin Echo MRSI acquisition without outer volume suppression was implemented, with the slice positioned along the anterior and posterior commissural planes (AC-PC) and placed just superior to the ventricles. At the GE-site (ACH), initial whole-slice MRSI acquisitions indicated that extracranial lipid suppression via cubical localization was needed to improve spectral quality. Therefore, the MRSI protocol was modified to include a semi-Localization by Adiabatic Selective Refocusing (sLASER) module. The rectangular sLASER boundary was realized using 10 kHz bandwidth GOIA-WURST Radiofrequency pulses (Deelchand et al., 2021) to minimize chemical shift displacement errors, and is prescribed to be as large as possible in-plane without extending into the extra-cranial tissue for each participant. At the Siemens sites with whole-slice MRSI, lipid contamination is removed in post-processing using established MRSI lipid removal algorithms (Bilgic et al., 2014; Tsai et al., 2019).

Technical innovations in *k*-space sampling strategies have enabled substantial reductions in scan time (Bogner et al., 2021). To ensure feasibility for pediatric imaging, a non-Cartesian, rosette *k*-space sampling trajectory (Li et al., 2015; Noll, 1997; Schirda et al., 2009; Senthil et al., 2023) was implemented to acquire 7.5 mm in plane multi-voxel spectroscopic data in 4-5 minutes, thus minimizing participant burden and motion. Automated high-order shimming was implemented at all sites to reduce magnetic field (B_0_) inhomogeneities and boost signal retention during data acquisition (Xu et al., 2023).

## 4 Data Organization, Processing, and Sharing

The MRI acquisition framework described here forms part of a broader infrastructure developed within the C-PIP platform to support standardized automated data organization processing and sharing in Canadian pediatric hospital imaging centers. Imaging data acquired at each site are transferred to a centralized data management environment where datasets are organized according to standardized conventions and integrated with phenotypic data (e.g., demographic, anthropometric, and cognitive measures) collected within the platform. Imaging data are organized using standardized data structures (Brain Imaging Data Structure [BIDS]) (Gorgolewski et al., 2016; Poldrack et al., 2024) to facilitate automated processing pipelines and cross-site reproducibility. Centralized preprocessing pipelines are applied to structural, diffusion, and functional imaging data to ensure consistent versioning, data handling, and quality control across sites. These pipelines incorporate widely used neuroimaging software tools and include procedures for artifact correction, motion assessment, and quality monitoring.

To support open science and collaborative research, the C-PIP platform is designed to enable secure data sharing and controlled access to curated datasets. Deidentified raw and processed imaging data are made available through a federated data sharing infrastructure following appropriate ethical approvals and data governance procedures. The C-PIP infrastructure adheres to FAIR-2 (Findable, Accessible, Interoperable, Reuseable) principles (Wilkinson et al., 2016), thus offering a sustainable resource for advancing the landscape of neuroscientific research. Detailed descriptions of the data infrastructure, processing pipelines, and data-sharing framework are provided elsewhere [Spinney et al. under review].

## 5. Travelling Subject Stability Study

### 5.1 Rationale

Larger sample sizes afforded by multisite neuroimaging datasets allow for increases in statistical power and sensitivity to detect reliable effects (Marek et al., 2022). However, this presents unique challenges for differentiating true biological effects from site- and system-level effects due to differences in scanner hardware and acquisition parameters. In this study, we investigated the validity of acquisition standardization in the C-PIP imaging protocol. We employed a ‘travelling participant’ approach, where human volunteers were scanned twice at each site to assess within- and between-site stability. Here, we focus specifically on the core sequences (structural T1w imaging, diffusion-weighted imaging, and multi-echo functional imaging) given their high sensitivity to scanner-related differences and use across the neurodevelopmental literature.

### 5.2 Study design

MRI data were acquired from three young adult human volunteers (age-range: 30-33 years, two females) between September – November 2025. All participants were healthy and had no MRI contraindications. Participants were scanned twice at each site, resulting in a total of six scans per participant (total: 18 individual scans). Participants were scanned with the full Child and Adolescent C-PIP protocol, which included the following core acquisitions analyzed here: (1) structural T1w; (2) multi-shell DWI; (3) two movie ME-fMRI runs (Partly Cloudy and La Luna). Participants were taken out of the scanner after their first session and repositioned for the second session following a short break (∼20 mins). At each site, the total scan duration for each participant across the two scanning sessions was approximately 2 hours. A flowchart of the study design is presented in Figure 2. Study procedures were conducted in accordance with the ethical principles stipulated in the World Medical Association (WMA) Declaration of Helsinki. All travelling participant volunteers provided written informed consent at each site prior to scanning.

**Figure 2.**
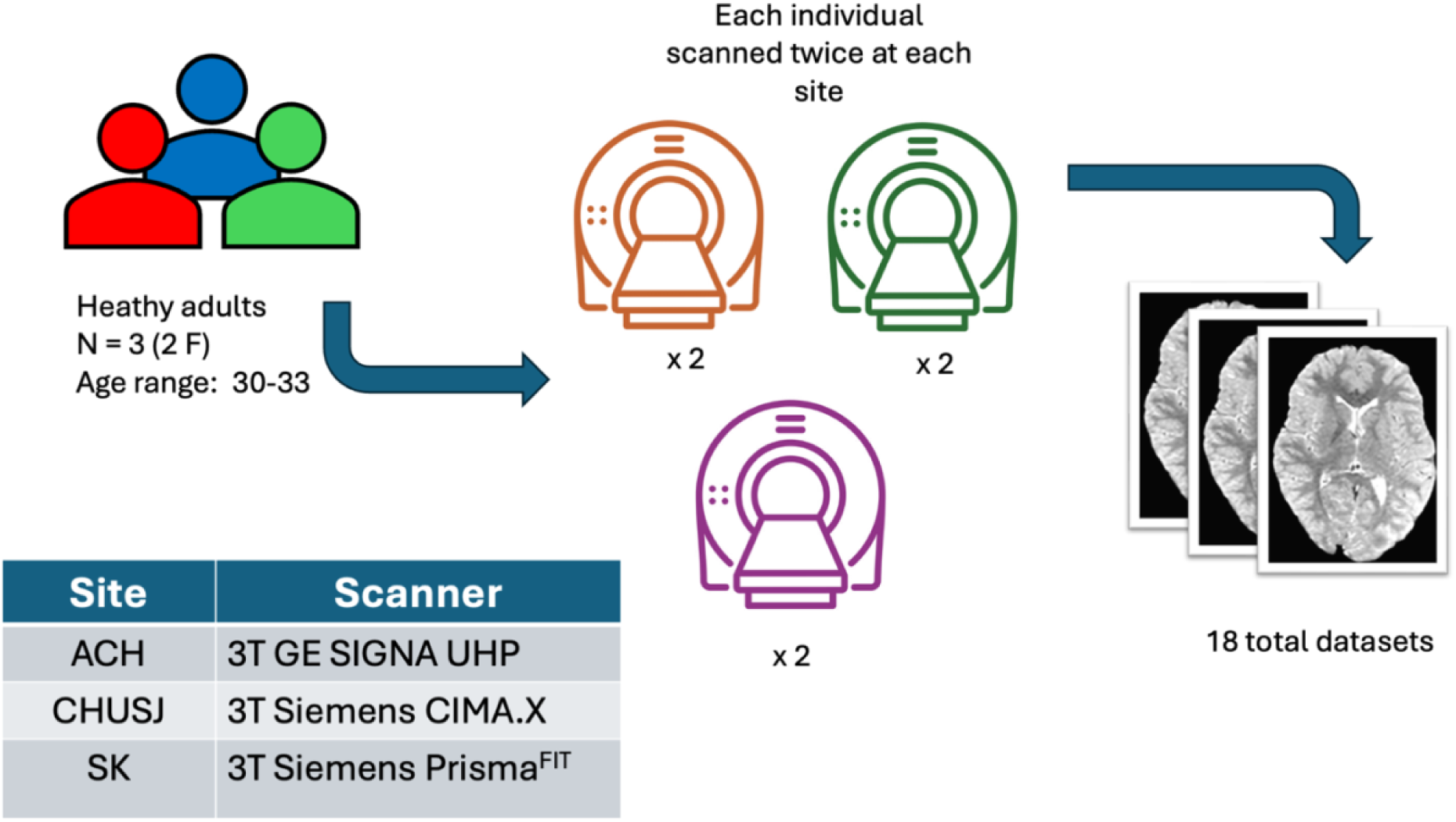
Study design for the C-PIP travelling subject stability study.

### 5.3 Image Processing Structural MRI

T1w images for each travelling subject were processed for each site using FreeSurfer (version 7.3.2, https://surfer.nmr.mgh.harvard.edu/), which consisted of several processing steps including skull stripping, Talairach transformation, surface extraction, spherical registration and parcellation of cortical and subcortical structures (Fischl, 2012). Given that each travelling subject had test-retest data, the longitudinal stream was chosen, whereby scans from each session were processed independently and then resampled to an unbiased within-subject template (Reuter et al., 2010). Cortical regions (62 regions-of-interest; 31 per hemisphere) were parcellated from the processed T1w images using the Desikan-Killiany (DK) atlas (Desikan et al., 2006), while subcortical structures (18 regions; 9 per hemisphere) were segmented using the Aseg atlas (Fischl et al., 2002). Region-specific averaged measures of cortical surface area (SA), cortical thickness (CT), cortical gray matter volume (vol), and subcortical gray matter volume (subvol) were computed from each participant’s scan.

#### Diffusion MRI

Diffusion images were processed centrally in MRtrix3 (version 3.0.5) (Tournier et al., 2019). To ensure comparability with prior DTI-based literature, the *b* = 1000 s/mm^2^ shell (30 directions) was extracted from each dataset and used for downstream processing. Processing steps followed the recommended steps in MRtrix3, which included (i) Denoising via Marchenko-Pastur Principal Component Analysis (Veraart et al., 2016), (ii) Gibbs ringing correction (Kellner et al., 2016), and (iii) eddy-current, motion, and susceptibility distortion correction (Andersson et al., 2017, 2016, 2003), with the latter using a separately collected reverse-phase encoded *b* = 0 series (PA direction). The preprocessed diffusion data were then up-sampled to a voxel size of 1 mm^3^ to improve downstream model-fitting and fitted using the tensor model (Basser et al., 1994) to estimate the diffusion signal within white matter voxels. Tensor maps from each subject were then visually assessed to ensure tensors were correctly oriented along the principal diffusion directions and aligned with known major white matter structures (i.e., corpus callosum, corticospinal tracts). Whole-brain metric maps of fractional anisotropy (FA) and mean diffusivity (MD) were computed to derive estimates of white matter microstructure.

Tract of interest analysis was performed using the John Hopkins University White Matter Atlas, (Hua et al., 2008). The following major white matter tracts of interest were chosen: (1) left and right Anterior thalamic radiation (ATR); (2) left and right Corticospinal tracts (CST), (3) Forceps major and minor (F_maj_; F_min_); (4) left and right Cingulum (CG), (5) left and right Inferior fronto-occipital fasciculus (IFO), (6) left and right Inferior longitudinal fasciculus (ILF), (7) left and right Superior longitudinal fasciculus (SLF), and (8) left and right Uncinate fasciculus (UF). This resulted in 16 individual white matter bundles. All tracts were warped from standard JHU-DTI stereotaxic coordinate space to each participant’s native diffusion space using linear and nonlinear registration in FSL and then binarized to create voxel masks. Lastly, for each participant, FA and MD values were averaged across all voxels within each tract mask to obtain tract-wise estimates of white matter microstructure.

#### Multi-echo fMRI

Multi-echo fMRI data for each run (Partly Cloudy and La Luna) were preprocessed using fMRIPrep (version 25.2.3) (Esteban et al., 2019). As an initial step, slice-time correction was applied to all echoes to align all slices with the midpoint of the TR (TR = 2.209 seconds; midpoint = 1.1045 seconds). Subsequent preprocessing steps were based on the first echo and included (1) motion correction using a reference BOLD image, and (2) susceptibility distortion correction via Phase Encoding Polarity (PEPOLAR) using a pair of reverse-phase encoded (AP-PA) spin-echo volumes acquired separately during the scan. The resulting spatial transformations were subsequently applied to the later echoes to preserve voxel-wise correspondence across all volumes in the fMRI series. Despite 10 dummy volumes being discarded from data acquired at CHUSJ and SK, the first 8 volumes were ignored from data acquired at all sites during processing to ensure to ensure temporal alignment across all sites. As per the default workflow for fMRIPrep, motion (rotations, translations, framewise displacement), physiological signal properties (cerebrospinal fluid, white matter, global signal) and their derivatives, were computed for each volume and outputted to be used as nuisance covariates in subsequent confound regression.

Processed data in subject-space were then passed into TEDANA (version 25.1.1), where each echo was optimally combined using the T2* combination method (DuPre et al., 2021; Kundu et al., 2017). At this stage, the 8 volumes that were ignored during fMRIprep processing were explicitly removed from the optimally combined data. BOLD and non-BOLD components were then estimated using a combination of principal (PCA) and independent component analyses (ICA) based on the Kundu decision tree (Kundu et al., 2012). Specifically, PCA was performed to decompose the optimally combined data to a fixed set of components to be included in ICA denoising. The number of components specified for PCA was chosen to be approximately half the number of volumes in each run. Following this, ICA was repeatedly run to identify the most stable components across a maximum number of 50 iterations using the “robustica” flag.

External regressors outputted from fMRIPrep, which include motion (rotations, translations, framewise displacement), physiological signal properties (cerebrospinal fluid, white matter, global signal), and their derivatives, were supplied during denoising to regress out nuisance variance and improve component classification. Classified components were then visually assessed to ensure that all BOLD-like components were correctly identified and accepted.

Confound regression was then performed on the optimally combined data using the same regressors used for ICA classification. Additionally, components identified and manually verified during ICA as “non-BOLD” (i.e., “rejected” components) were also included in the regression model as nuisance covariates. A high-pass filter was also applied during regression to attenuate low-frequency signals below a threshold of 0.01Hz. The data was then spatially standardized to a common space (Montreal Neurological Institute [MNI]) using the default ICBM 2009c Nonlinear Asymmetric template and each run was mean-centered and concatenated into a single timeseries. Lastly, using framewise displacement values estimated during fMRIprep processing (calculated using the Power algorithm) (Power et al., 2012), motion censoring was performed on the concatenated data to remove volumes exceeding a threshold of >0.2 mm. Table S1 in the Supplementary Materials provides a breakdown of how many volumes were removed for each participant.

### 5.4 Stability Analysis

#### Structural T1w and Diffusion MRI

For metrics derived from structural and diffusion MRI scans, test-retest stability was assessed independently at each site by comparing measurements obtained from repeated scanning sessions for the same participant. Stability was quantified using the absolute percentage difference between sessions, a metric that reflects the magnitude of variability between two repeated observations relative to their average value, ranging from 0% to 100%. Lower absolute percentage differences were interpreted as evidence of higher test-retest stability.

Cross-site agreement was evaluated by comparing measurements acquired at different sites. Percentage differences were calculated for each pair of site-specific measurements for the same individual (CHUSJ vs. ACH, SK vs. ACH, SK vs. CHUSJ), yielding values potentially ranging from −100% to 100%. Within-site values for each participant were averaged prior to calculating cross-site percentage differences to minimize between-session noise. Cross-site percentage difference values near 0% indicate strong measurement agreement between sites and minimal influence of site-related factors. Positive or negative values indicate systematic differences between sites, with the sign reflecting the direction of the bias in favor of a specific site.

#### Multi-echo fMRI

***Functional connectivity analysis.*** Region-to-region functional connectivity matrices were generated for each participant by computing Pearson correlations between time series averaged across cortical regions defined from 200 regions-of-interest (ROI) in the Schaefer 200 atlas (Schaefer et al., 2018). Edgewise functional connectivity between ROIs from the same individual scanned at different sites were correlated to assess similarity in connectivity profiles across sites, while site-specific inter-subject similarity was assessed by correlating functional connectomes from all participants scanned at the same site to assess whether functional connectivity profiles differed systematically between sites.

***ROI-based Test-Retest Video Viewing Correlations.*** In addition to estimating edgewise functional connectivity, ROI-based inter-scan correlation analysis was conducted on the concatenated data to assess whether watching the same visual stimuli during scanning elicited repeatable response patterns within participants, within and across sites (Nastase et al., 2019). Within-person within-site consistency of activation patterns was evaluated by first correlating ROI-to-ROI preprocessed timeseries between session pairs for each participant separately, then averaging across the sample to produce a combined site-specific correlation map (n = 3 participants). Across-site consistency was assessed by correlating all unique session-wise ROI timeseries from scans acquired from the same individual at different sites (ACH vs. CHUSJ, ACH vs. SK, CHUSJ vs. SK). These were then averaged to produce a combined within-subject, across-site map for each pairwise site comparison (12 unique pairings per site-pair, as each person was scanned twice at each site). Higher positive correlations indicate greater similarity in activation patterns.

### 5.5 Results Structural T1w

Region-wise structural measures demonstrated high test-retest stability with all values under 15% (Figure 3A). For ACH, values ranged from 0.0%-14.0% for surface area (SA), 0.0%-7.1% for cortical thickness (CT), 0.0%-13.9% for volume, and 0.0%-13.3% for subcortical volume. CHUSJ showed lower overall ranges, with SA ranging from 0.0%-12.7%, CT from 0.0%-6.6%, volume from 0.0%-10.6%, and subcortical volume from 0.0%-8.9%. SK exhibited the lowest variability, with SA ranging from 0.0%-4.7%, CT from 0.0%-5.6%, volume from 0.0%-6.0%, and subcortical volume from 0.0%-8.7%. Common regional patterns of test-retest stability emerged across sites, with the entorhinal cortex (maximum: 14.0%) and accumbens (maximum: 12.6%) showing the greatest between-session variability (Supplementary Figure 9).

**Figure 3.**
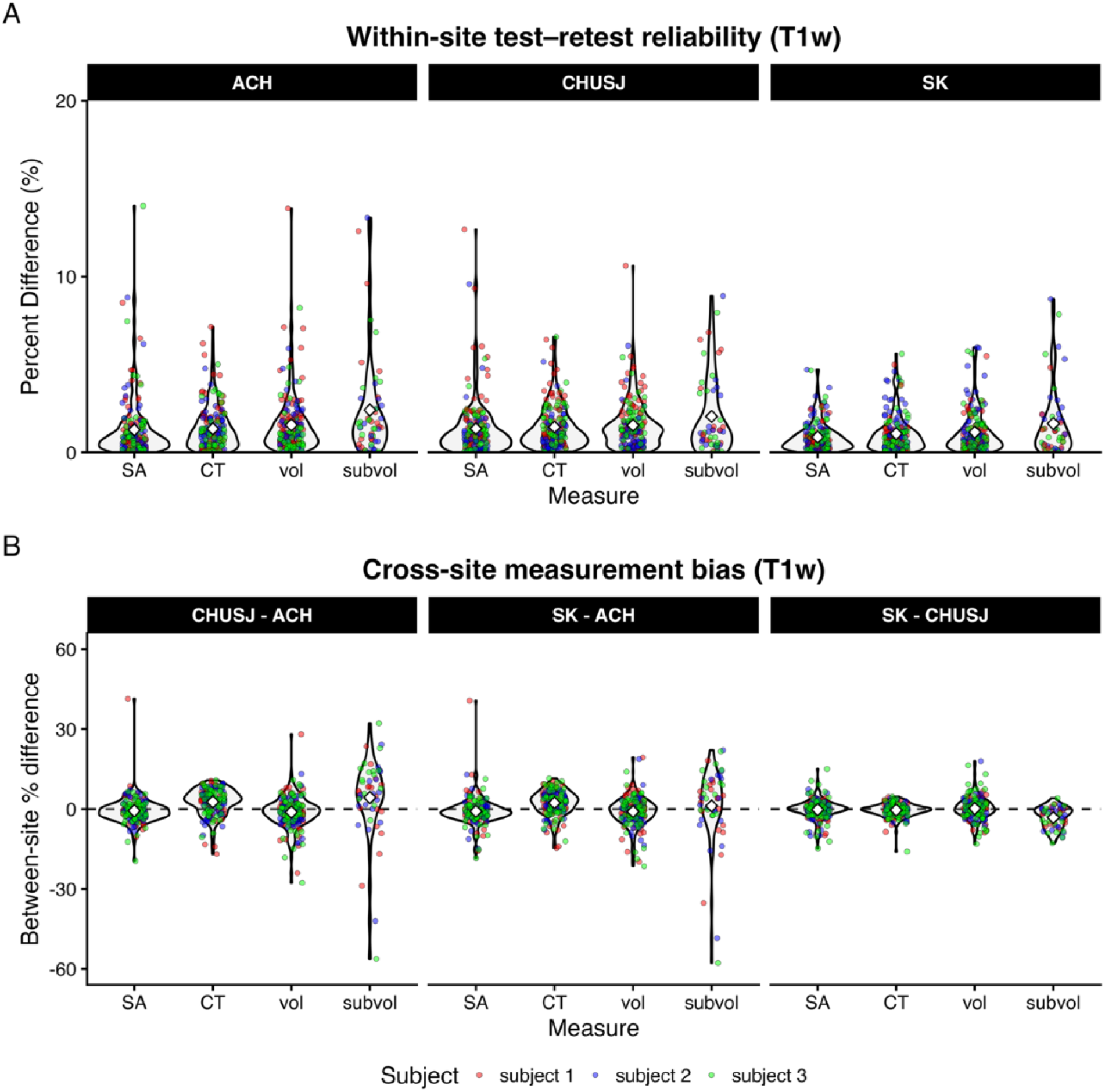
Stability analyses of structural (T1w) metrics. **(A)** Absolute percentage differences between repeated sessions within each site. **(B)** Cross-site comparisons between measurements acquired between sites (CHUSJ vs ACH; SK vs ACH; SK vs CHUSJ). **CHUSJ vs ACH:** positive differences indicate CHUSJ has higher values. **SK vs ACH:** positive differences indicate SK has higher values. **SK vs CHUSJ:** positive differences indicate SK has higher values. **Abbreviations: SA** (surface area); **CT** (cortical thickness); **vol** (grey matter volume); **subvol** (subcortical volume); **ACH** (Alberta Children’s Hospital); **CHUSJ** (Centre hospitalier universitaire Sainte-Justine); **SK** (Hospital for Sick Children – SickKids). **Subject-1**: Red; **Subject-2**: Blue; **Subject-3**: Green. White diamonds denote the average value for that measure across subjects.

Measurement agreement was generally good across all pairwise between-site comparisons (Figure 3B). Similar to within-site measures, subcortical volume demonstrated the greatest variability in percentage difference values, particularly for measurements collected between ACH and CHUSJ (range: −56.2% - 32.2%), and between ACH and SK (range: −57.8% - 22.1%).

#### Diffusion MRI

Tract-averaged diffusion measures also demonstrated good test-retest stability, with low percent differences observed at each site (Figure 4A) for FA (ACH range: 0.0% - 3.9%; CHUSJ range: 0.0% - 6.4%; SK range: 0.2% - 3.0%), and MD (ACH range: 0.0% - 3.8%; CHUSJ range: 0.0% - 3.8%; SK range: 0.0% - 4.4%). Test-retest stability was broadly consistent across tracts (Supplementary Figure 10), with most mean percentage difference values clustered around 1% across all tracts for FA and MD, although slightly higher variability was observed in some association pathways, specifically, for FA in the left SLF (maximum 6.4%) and right UF (maximum 4.5%).

**Figure 4.**
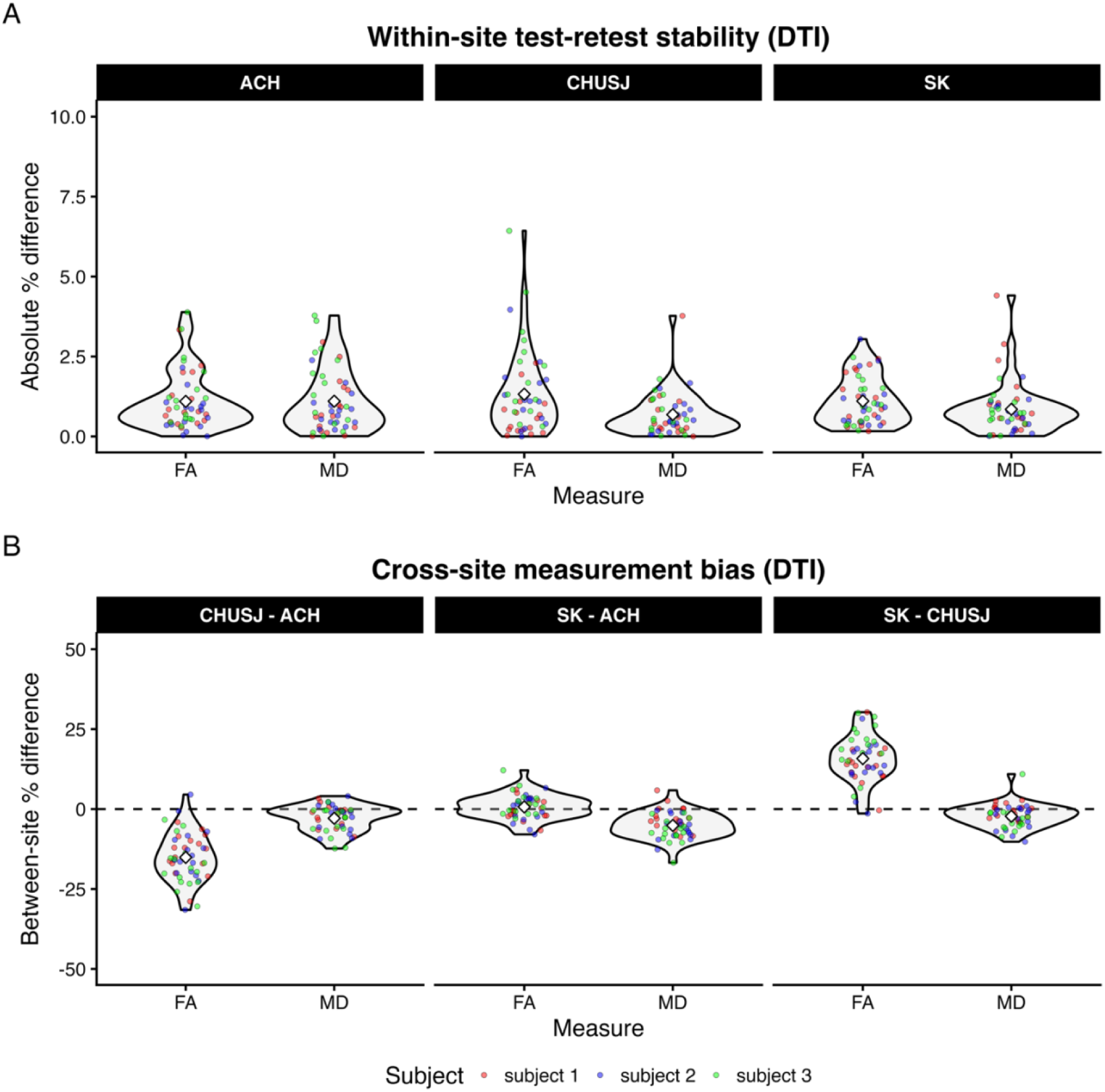
Stability analyses of diffusion MRI metrics. **(A)** Absolute percentage differences between repeated sessions within each site. **(B)** Cross-site comparisons between measurements acquired between sites (CHUSJ vs ACH; SK vs ACH; SK vs CHUSJ). **CHUSJ vs ACH:** positive differences indicate CHUSJ has higher values. **SK vs ACH:** positive differences indicate SK has higher values. **SK vs CHUSJ:** positive differences indicate SK has higher values. **Abbreviations: FA** (fractional anisotropy); **MD** (mean diffusivity); **ACH** (Alberta Children’s Hospital); **CHUSJ** (Centre hospitalier universitaire Sainte-Justine); **SK** (Hospital for Sick Children – SickKids). **Subject-1**: Red; **Subject-2**: Blue; **Subject-3**: Green. White diamonds denote the average value for that measure across subjects.

When looking at between-site measurement agreement (Figure 4B), CHUSJ demonstrated a site-related bias for lower tract-averaged FA compared to ACH (range: −31.5% - 4.5%) and SK (range: −1.4% - 30.3%).

#### Multi-echo fMRI

***Functional connectivity analysis.*** Figure 5 presents cross-correlation matrices of associations between session pairs of whole brain functional connectomes for the same participant scanned at different sites (panel A), and for all participants scanned at the same site (panel B). Results demonstrate that participant-specific connectivity patterns were preserved across sites, with mean functional connectivity across sites for the same subject being higher (mean Pearson’s *r*; subject-1: 0.79; subject-2: 0.76; subject-3: 0.81; Figure 5 panel A) compared to connectivity profiles across participants within the same site (mean Pearson’s *r*; ACH: 0.60; CHUSJ: 0.58; SK: 0.59; Figure 5 panel B), suggesting that inter-individual differences in functional brain organization are robust to site-related variation.

**Figure 5.**
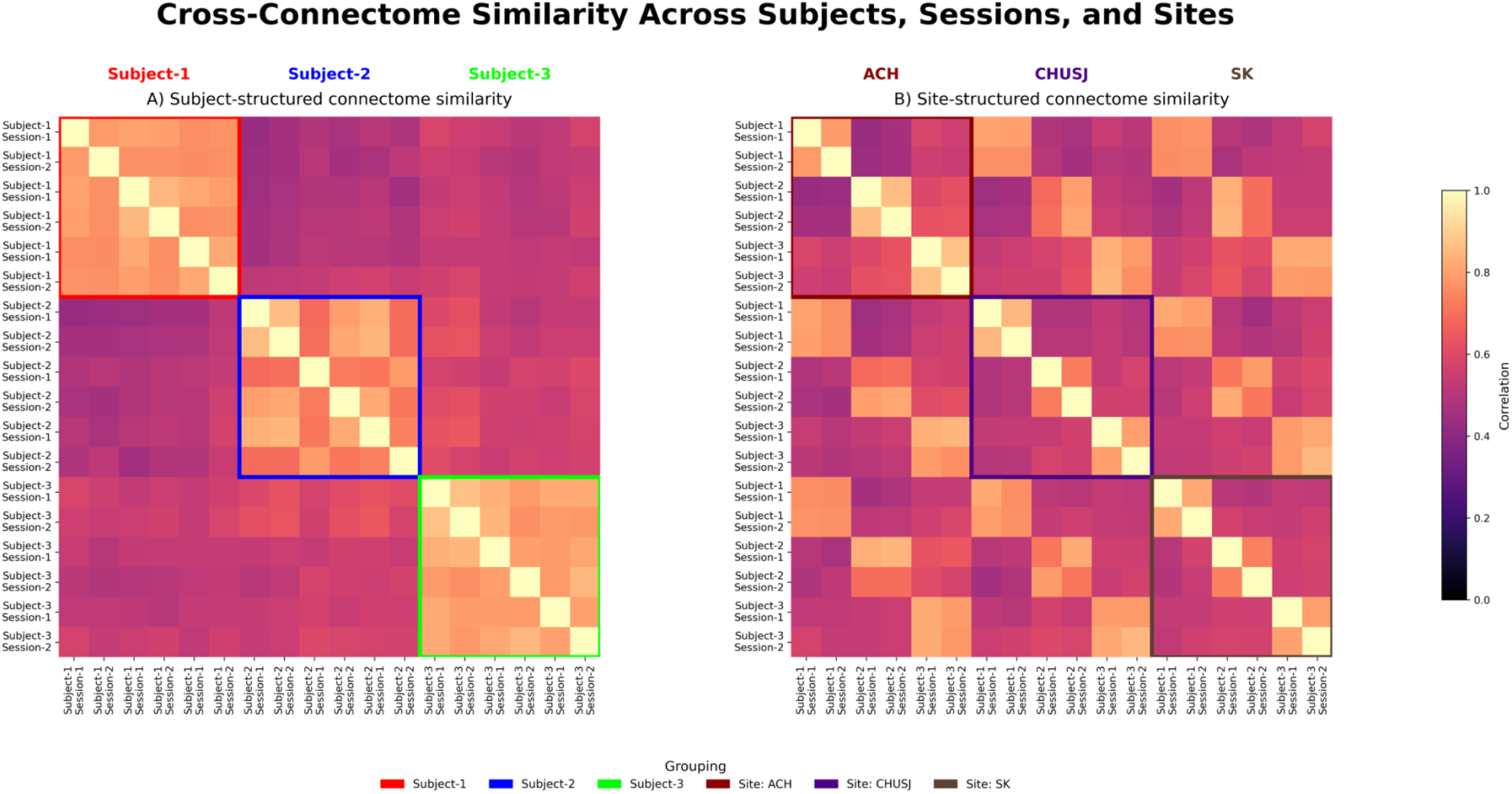
Stability analyses of multi-echo fMRI functional connectivity. Panel A-B: Heatmaps depicting within- and between-site associations of whole-brain functional connectomes generated from the same subject across sites (Panel A) and across different subjects within sites (Panel B). **Abbreviations: ACH** (Alberta Children’s Hospital); **CHUSJ** (Centre hospitalier universitaire Sainte-Justine); **SK** (Hospital for Sick Children – SickKids).

***ROI-based Test-Retest Video Viewing Correlations.*** This analysis showed similar spatial patterns of functional activation across participants within each site, with stronger test-retest correlations observed in the ventral visual cortices (Figure 6, blue panels), although some variability was evident. For between-site analysis, activation patterns for the same individual across sites also demonstrated reasonable agreement (Figure 6, orange panels), indicating that despite site-related differences in scanning systems, video-viewing evoked similar patterns of activity.

**Figure 6.**
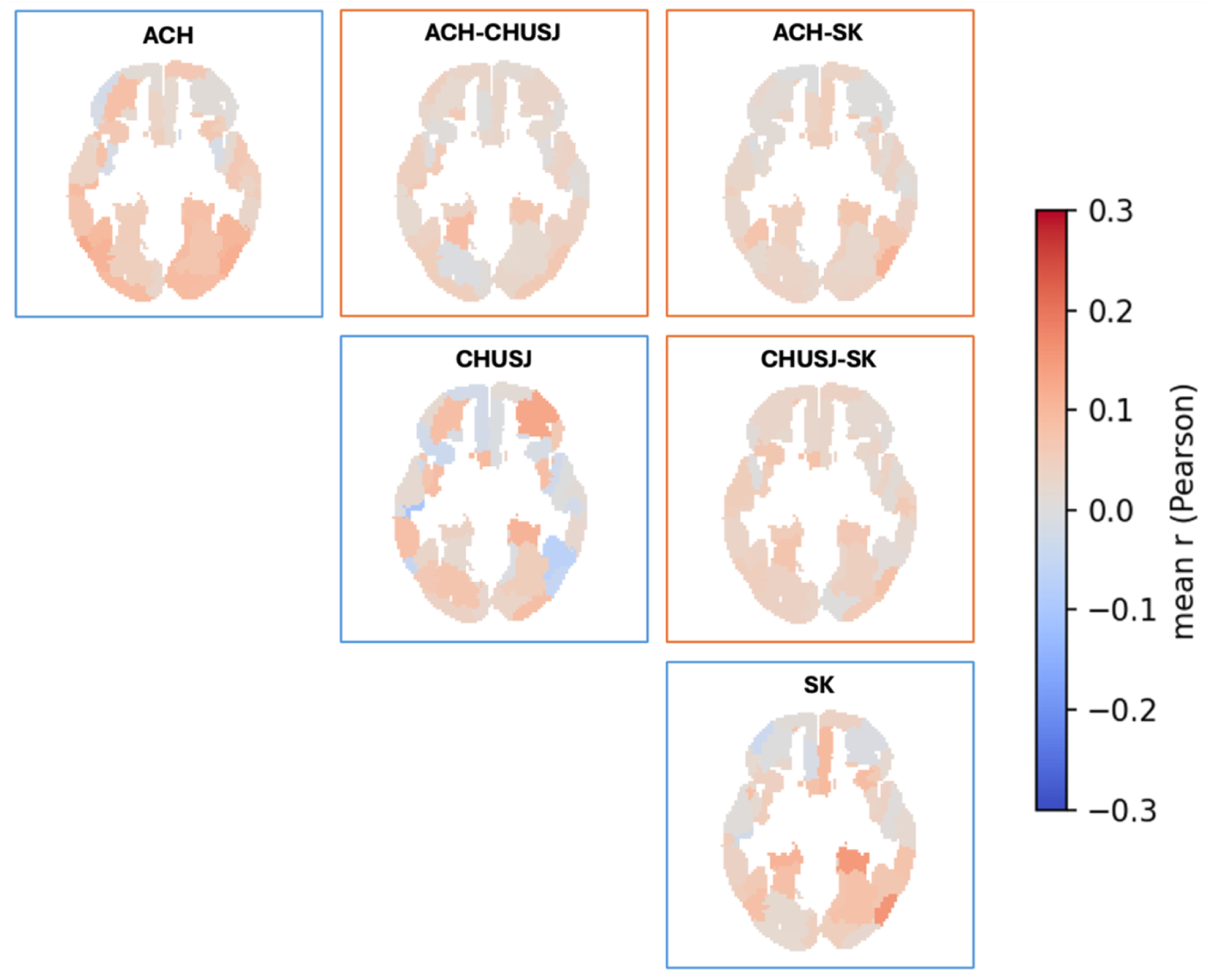
Within-subject test-retest regional BOLD video-viewing timeseries correlations, within-site (N=3 per site; diagonal blue panels) between-sites (N=12 pairs averaged, 3 participants and 2 acquisitions per site; orange panels). Maps were generated by averaging all unique pairings and visualized in the axial plane (slice z = 30). **Abbreviations: ACH** (Alberta Children’s Hospital); **CHUSJ** (Centre hospitalier universitaire Sainte-Justine); **SK** (Hospital for Sick Children – SickKids).

## 5. Discussion

A central objective of the C-PIP MRI protocol was to characterize normative patterns of brain development in a Canadian-specific reference cohort spanning infancy to late adolescence. C-PIP has achieved harmonized data acquisition across multiple pediatric imaging centres while accommodating differences in scanner hardware and vendor-specific implementations. Rather than enforcing identical acquisition parameters, the protocol emphasized alignment of key imaging characteristics, including spatial resolution, diffusion sampling, acquisition geometry, and image contrast. Results from the travelling-participant study suggest that this strategy was largely successful. Structural, diffusion, and functional imaging measures demonstrated strong within-site stability and generally adequate agreement across sites, supporting the feasibility of pooling data across centres for developmental neuroimaging research. Some systematic site-related differences were observed (i.e., lower FA at CHUSJ compared to the other two sites) that can be further addressed through established statistical harmonization approaches during analysis (Hu et al., 2023; Jodoin et al., 2025).

Several features distinguish the C-PIP imaging framework from existing developmental neuroimaging initiatives. First, the platform spans a broad brain developmental range from infancy through adolescence using harmonized multimodal imaging acquired across three pediatric research-dedicated MRI systems. Second, by recruiting participants in three different provinces, the platform provides a uniquely Canadian resource for studying brain development within geographically diverse populations while facilitating multisite collaboration. Third, imaging data are integrated with harmonized processing pipelines and data-management infrastructure designed to support reproducible analyses and data sharing. Finally, in addition to the commonly used sequences in the developmental literature (i.e., structural, diffusional, and functional imaging), the inclusion of advanced quantitative imaging modalities, such as ihMT, STAGE, and novel acquisition strategies (i.e., MRSI), extends beyond conventional imaging, creating opportunities for the development and validation of novel neuroimaging biomarkers (Saltarelli et al., 2025).

An important feature of the C-PIP framework is the inclusion of a dedicated Infant and Toddler protocol. Imaging during the first years of life presents unique challenges related to rapidly changing tissue contrast, limited scan tolerance, and safety considerations associated with natural-sleep imaging (Copeland et al., 2021; Spann et al., 2023). The modified protocol incorporates age-appropriate familiarisation strategies, structural imaging contrasts, shortened acquisitions, acoustic considerations, and sequence-specific adaptations to accommodate SAR constraints while maintaining a common multimodal imaging framework across age groups. These adaptations enable the collection of comparable imaging measures across a developmental period that remains underrepresented in large-scale neuroimaging studies.

Researchers using C-PIP imaging data should nevertheless consider site-related variation when designing analyses. Inclusion of scanner site as a covariate, application of harmonization methods such as ComBat (Bayer et al., 2022; Orlhac et al., 2022), and use of centralized processing pipelines are recommended where appropriate. Investigators should also consider developmental stage when interpreting imaging measures, particularly for modalities whose biological interpretation may vary across age.

Despite demonstrating adequate test-retest and cross-site reliability for the C-PIP core sequences, the travelling-participant study included only three participants and was conducted in healthy adults rather than children. While this design provides a practical assessment of scanner-related variability independent of developmental factors, it does not fully capture challenges associated with pediatric imaging, including developmental differences in tissue properties and motion. Future work will evaluate longitudinal stability of imaging measures within the C-PIP cohort and further refine harmonization strategies as the platform continues to grow.

## Conclusion

The C-PIP MRI protocol establishes a harmonized multisite framework for pediatric neuroimaging across three Canadian pediatric research centers. Through coordinated acquisition of structural, diffusion, functional, and advanced quantitative MRI measures, the protocol supports comprehensive characterization of brain development from infancy through adolescence. Travelling-participant analyses demonstrate strong within-site stability and acceptable cross-site reproducibility for core imaging measures, supporting the use of pooled multisite datasets. Together with centralized processing and data-sharing infrastructure, the C-PIP platform provides a valuable resource for investigating normative neurodevelopment, identifying developmental disruptions, and enabling future collaborative studies of pediatric neurological and psychiatric disorders.

## CRediT author statement

**Mervyn Singh:** Conceptualization, Writing - Original Draft, Writing - Review & Editing, Methodology, Formal analysis, Investigation, Visualization. **Shireen Parimoo:** Methodology, Investigation, Data Curation, Writing - Review & Editing. **Vicente Enguix:** Methodology, Investigation, Writing - Review & Editing. **Filomeno Cortese:** Methodology, Project administration, Investigation, Writing - Review & Editing. **Milton Camacho:** Methodology, Software, Data Curation, Writing - Review & Editing. **Sean Spinney:** Methodology, Software, Data Curation, Writing - Review & Editing. **Samantha J. Lynch:** Writing - Review & Editing. **Helen L. Carlson:** Methodology, Writing - Review & Editing. **Lara M. Leijser:** Methodology, Writing - Review & Editing. **Christine L. Tardif:** Methodology, Writing - Review & Editing**. Ilana R. Leppert:** Methodology, Writing - Review & Editing. **G. Bruce Pike:** Methodology, Writing - Review & Editing. **Emile Kadalie:** Methodology, Visualization, Writing - Review & Editing. **Jamie Near:** Methodology, Writing - Review & Editing. **Chatura Kumaragamage:** Methodology, Writing - Review & Editing. **Riya Chitroda:** Formal Analysis, Writing - Review & Editing. **Michel Louis Lauzon:** Methodology, Writing - Review & Editing. **Kathryn Y. Manning:** Writing - Review & Editing. **Gopal Varma:** Methodology, Writing - Review & Editing. **Tomas Paus:** Methodology, Writing - Review & Editing. **Ashley D. Harris:** Methodology, Writing - Review & Editing. **Gregory A. Lodygensky:** Methodology, Writing - Review & Editing. **Miriam H. Beauchamp:** Writing - Review & Editing. **Donald J. Mabbott:** Writing - Review & Editing. **Daniel C. Kopala-Sibley:** Writing - Review & Editing. **Jacob A. S. Vorstman:** Writing - Review & Editing. **Keith Owen Yeates:** Writing - Review & Editing. **Paul D. Arnold:** Writing - Review & Editing. **Sébastien Jacquemont:** Writing - Review & Editing. **Darren S. Kadis:** Project administration, Methodology, Funding acquisition, Writing - Review & Editing. **Sarah Lippé:** Project administration, Funding acquisition, Writing - Review & Editing. **Lune Bellec:** Software, Project administration, Methodology, Funding acquisition, Writing - Review & Editing. **Catherine Lebel:** Supervision, Conceptualization, Project administration, Funding acquisition, Methodology, Writing - Review & Editing. **Patricia Conrod:** Conceptualization, Writing - Review & Editing, Resources, Project administration, Funding acquisition. **Signe Bray:** Supervision, Conceptualization, Writing - Review & Editing, Methodology, Resources, Project administration, Funding acquisition. **Anne L. Wheeler:** Supervision, Conceptualization, Writing - Original Draft, Writing - Review & Editing, Methodology, Resources, Project administration, Funding acquisition.

## Declaration of Competing Interest

The authors declare the following financial interests/personal relationships which may be considered as potential competing interests: G. Varma is listed as a co-inventor on patents related to the ihMT technique. L. Bellec is a consultant for the ENIGMA consortium, funded through an NIH R01 grant. C. Lebel receives research support from GE Healthcare. J.A.S. Vorstman serves as a consultant for NoBias Therapeutics Inc. for the design of a clinical trial in children with the 22q11.2 deletion. S. Lippé serves as a consultant for Taysha Gene Therapies for the phenotypic analyses of a clinical trial in children with Rett Syndrome. These relationships did not influence the content of this manuscript. All other authors report no competing financial interests or personal relationships that would unduly influence the work reported in this manuscript.

## Acknowledgments

The Canadian Pediatric Imaging Platform (C-PIP) is proudly supported by a Platform grant from the Brain Canada Foundation through the Canada Brain Research Fund, with S. Bray being the Nominated Principal Applicant. Financial support is provided from Health Canada, and the Funding Partners: CHUSJ Foundation, CRACHUSJ Centre IMAGINE, SickKids Program in Neuroscience and Mental Health, Precision Child Health Partnership (PCHP), SickKids Garry Hurvitz Centre for Brain and Mental Health (GH-CBMH), SickKids Garron Family Cancer Centre, Alberta Children’s Hospital Research Institute (ACHRI), Alberta Children’s Hospital Foundation (ACHF), Hotchkiss Brain Institute (HBI), Owerko Centre for Neurodevelopment and Child Mental Health, Mathison Family Foundation. The funding bodies played no role in study design, data collection and analysis, decision to publish or preparation of the manuscript. The authors would like to declare the following funding sources: M. Singh is supported by a Harley Hotchkiss Samuel Weiss Postdoctoral Fellowship. S. J. Lynch is supported by a Canadian Institutes of Health Research (CIHR) Postdoctoral Fellowship (FRN194058). H. L. Carlson is funded by a grant awarded by the Heart and Stroke Foundation of Canada. G. B. Pike is funded by a CIHR Foundation grant and is a Campus Alberta Innovates Program Chair. R. Chitroda is supported by an Owerko Centre Graduate Scholarship. K.Y. Manning is funded by a Natural Sciences and Engineering Research Council of Canada (NSERC) Discovery grant. D.C. Kopala- Sibley is funded through a CIHR Project grant and an NSERC Discovery grant. A.D. Harris receives salary support through a Canada Research Chair position in MR spectroscopy in Brain injury and Pain, and is funded through grants awarded by CIHR, NSERC and The Arthritis Society (TAS). L. Bellec received funding from NSERC, IVADO, the Courtois foundation and was an FRQ-S senior scholar during part of this research. C. Lebel receives salary support through the Canada Research Chair position in Pediatric Neuroimaging. M.H. Beauchamp receives salary support through a Canada Research Chair position in Pediatric Traumatic Brain Injury and is funded through grants from CIHR. K. Owen Yeates is funded through a Network Catalyst grant awarded by CHIR and a grant awarded by the Canada Foundation of Innovation. J.A.S. Vorstman is supported by The SickKids Psychiatry Associates Chair in Developmental Psychopathology. P. D. Arnold is supported by the Cuthbertson and Fischer Chair in Pediatric Mental Health. S. Lippé receives salary support through a Canada Research Chair position in Pediatric Neurodiversity and is funded through grants from CIHR. A.L. Wheeler is funded through grants from CIHR. This research was enabled in part by support provided by Compute Ontario (computeontario.ca) and the Digital Research Alliance of Canada (alliancecan.ca).

