## Supplementary Materials for "The Canadian Pediatric Imaging Platform (C-PIP) Reference Sample: Neuroimaging Protocol and Cross-site Validation for Harmonized Multimodal Pediatric Imaging"

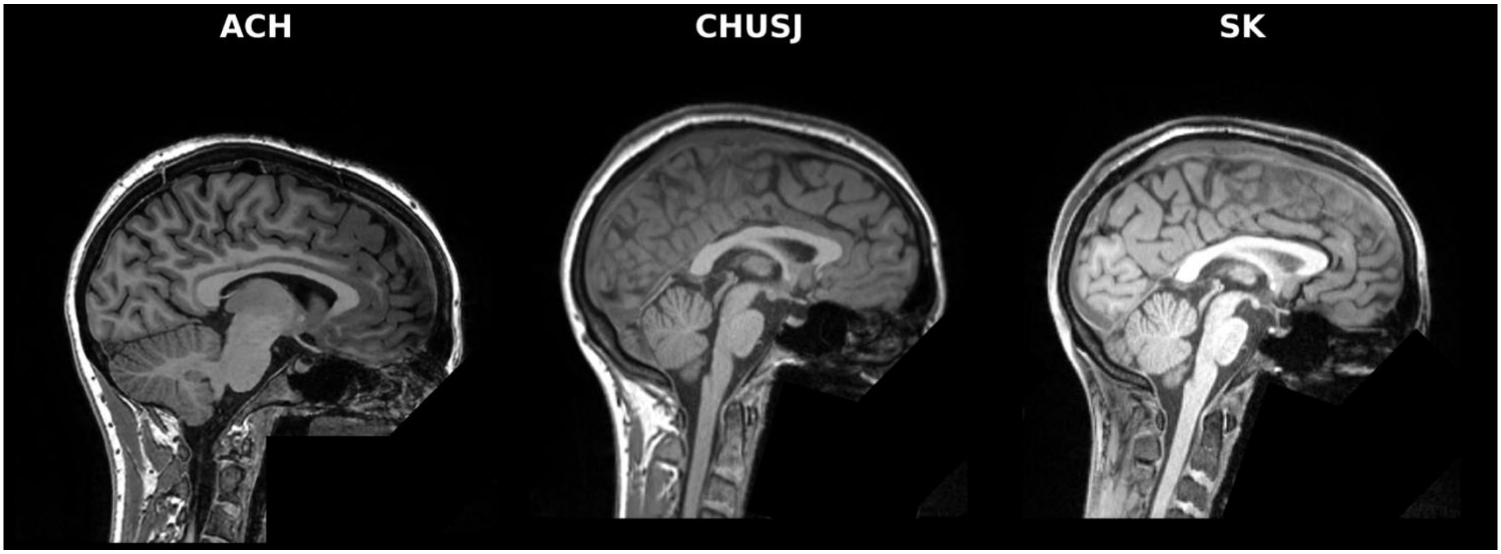


**Supplementary Figure 1.** Representative sagittal slices of T1-weighted images acquired from a single adult travelling participant (age 31) at each of the three C-PIP sites: (Left Panel) Alberta Children's Hospital (ACH); (Middle Panel) Centre hospitalier universitaire Sainte-Justine (CHUSJ); (Right Panel) Hospital for Sick Children – SickKids (SK). Raw images are shown. Data have been defaced to protect the identity of the participant.

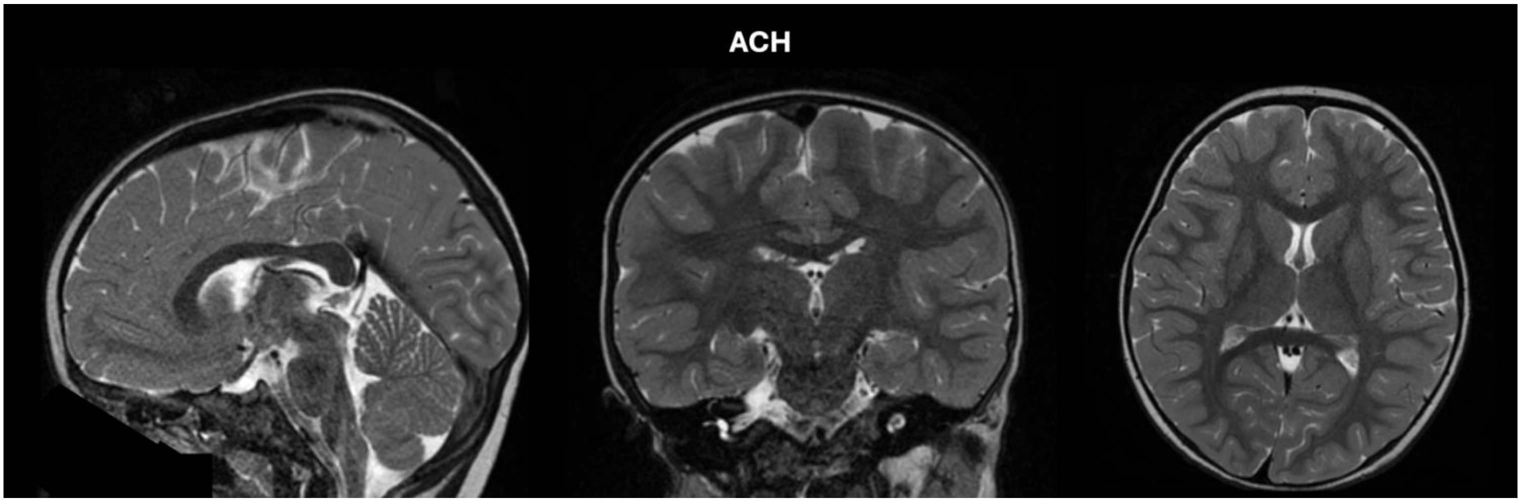


**Supplementary Figure 2.** Representative sagittal (left), coronal (middle) and axial (right) slices of T2-weighted images acquired from an infant participant at the Alberta Children's Hospital (ACH). Raw images are shown. Data have been defaced to protect the identity of the participant.

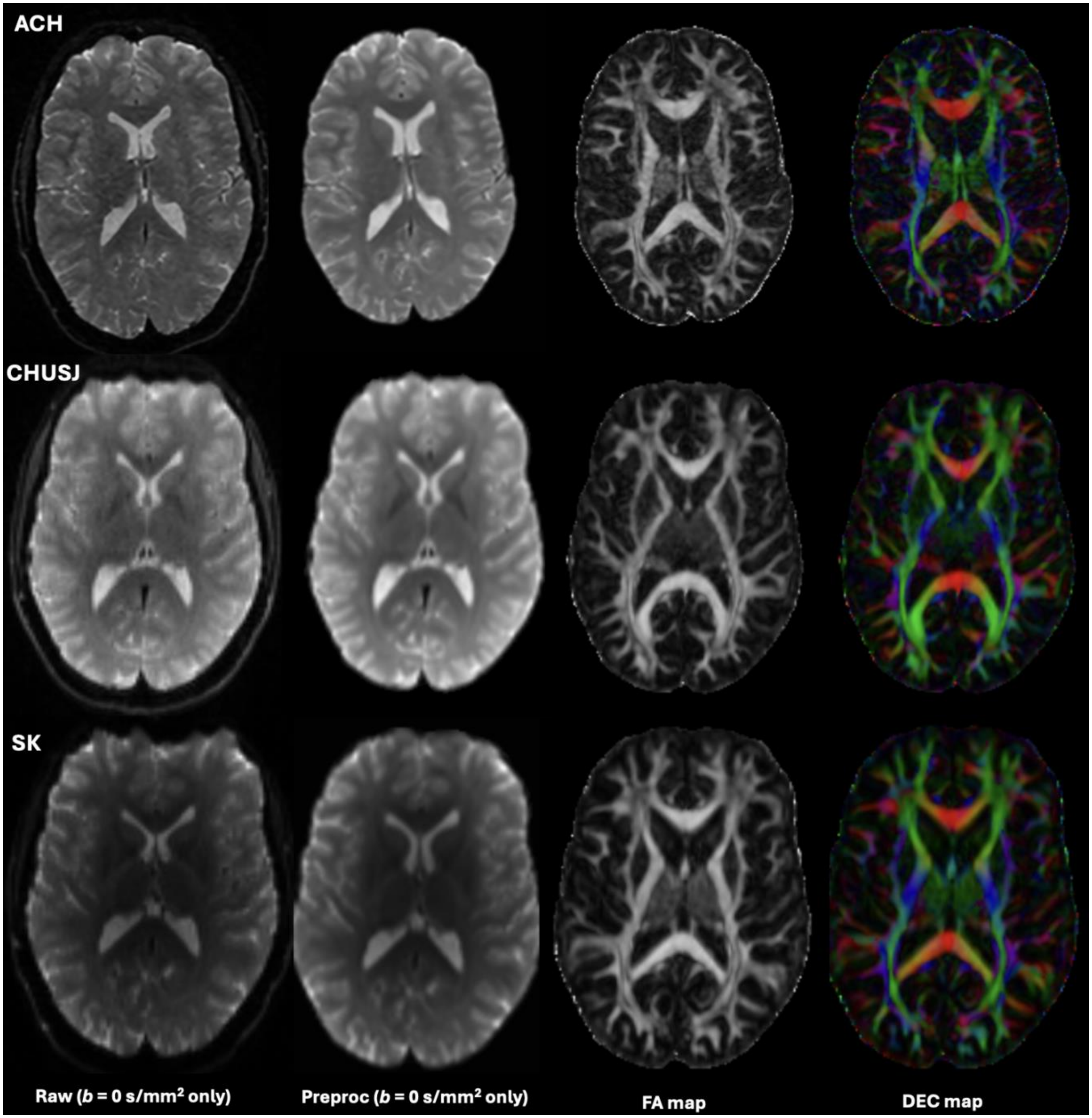


**Supplementary Figure 3.** Representative axial slices of diffusion MRI images acquired from a single adult travelling participant (age 31) at each of the three C-PIP sites: (Top panel) Alberta Children's Hospital (ACH); (Middle panel) Centre hospitalier universitaire Sainte-Justine (CHUSJ); (Bottom panel) Hospital for Sick Children – SickKids (SK). Raw, preprocessed and DTI-derived maps ($b$ = 1000 s/mm$^2$ only; fractional anisotropy [FA], directionally encoded color maps [DEC]) shown.

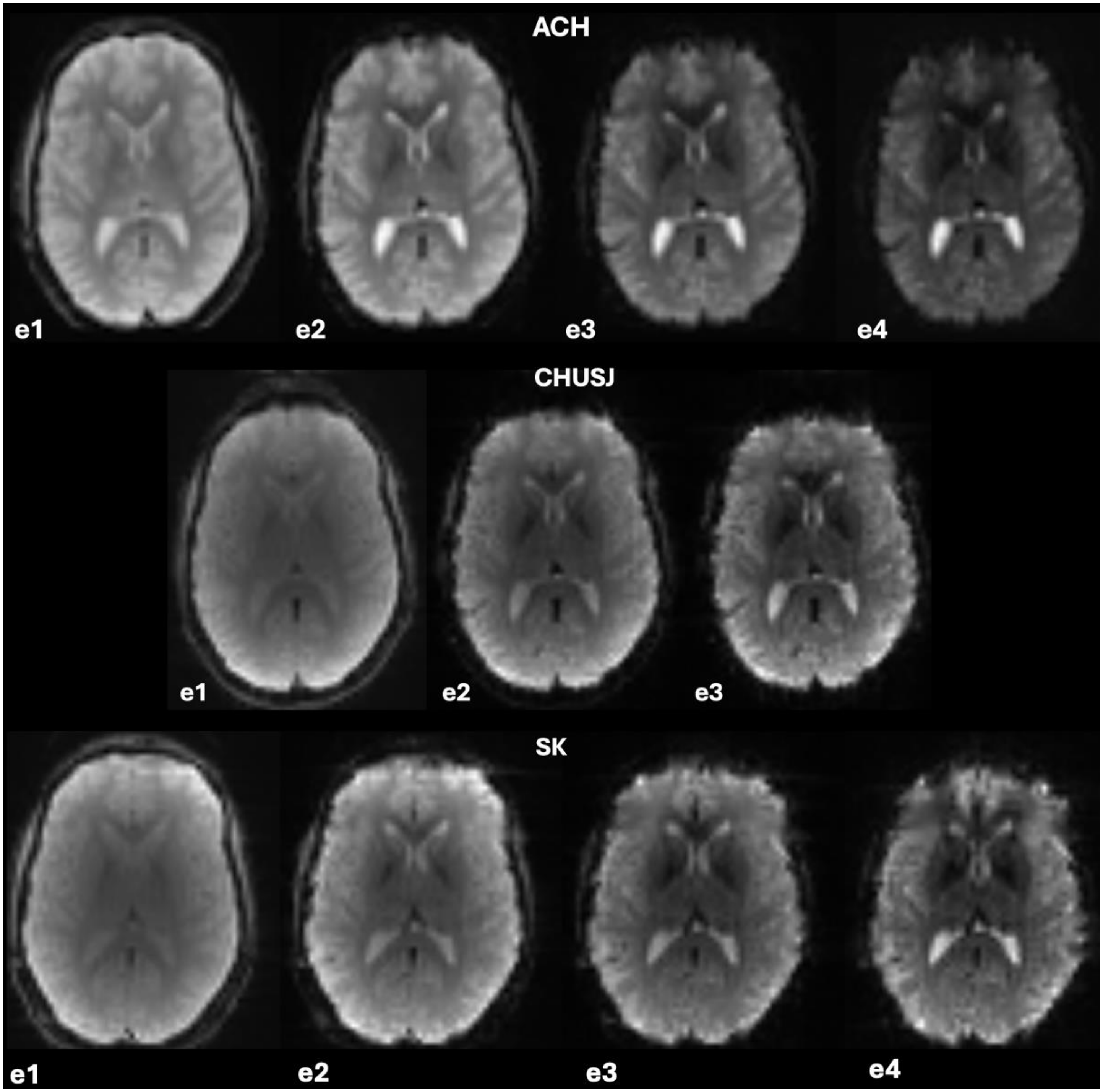


**Supplementary Figure 4.** Representative axial slices of raw Multi-echo fMRI images for each echo acquired from a single adult travelling participant (age 31) at each of the three C-PIP sites: (Top Panel) Alberta Children's Hospital (ACH); (Middle Panel) Centre hospitalier universitaire Sainte-Justine (CHUSJ); (Bottom Panel) Hospital for Sick Children – SickKids (SK). Abbreviations: e1-e4: echoes 1-4. Only the La Luna run is shown.

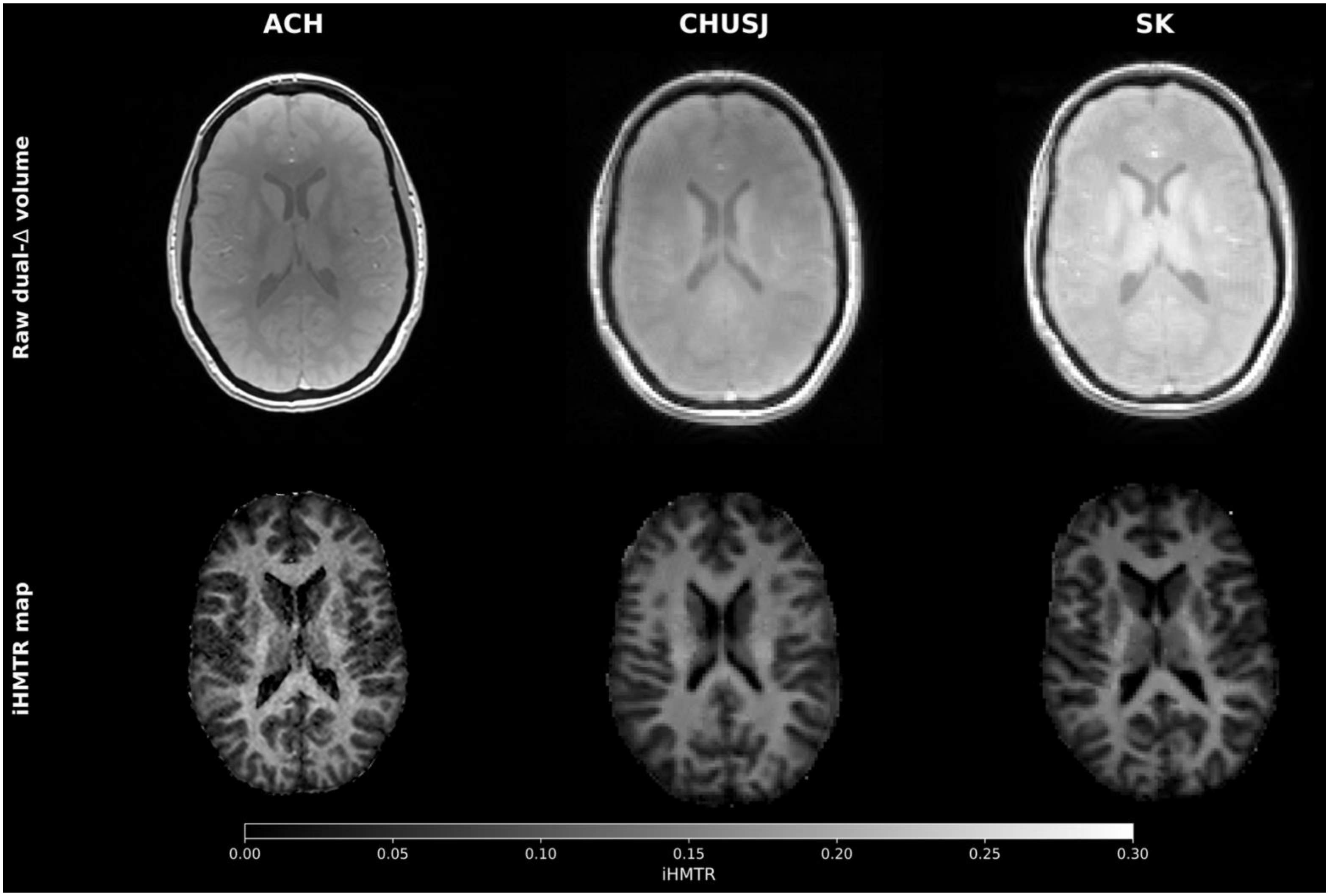


**Supplementary Figure 5.** Representative axial slices of inhomogeneous magnetization transfer imaging (ihMT) data acquired from a single adult travelling participant (age 31) at each of the three C-PIP sites: (Left panel) Alberta Children's Hospital (ACH); (Middle panel) Centre hospitalier universitaire Sainte-Justine (CHUSJ); (Right Panel) Hospital for Sick Children – SickKids (SK). Top row: raw dual MT preparation data; Bottom row: ihMT ratio (ihMTR) map.

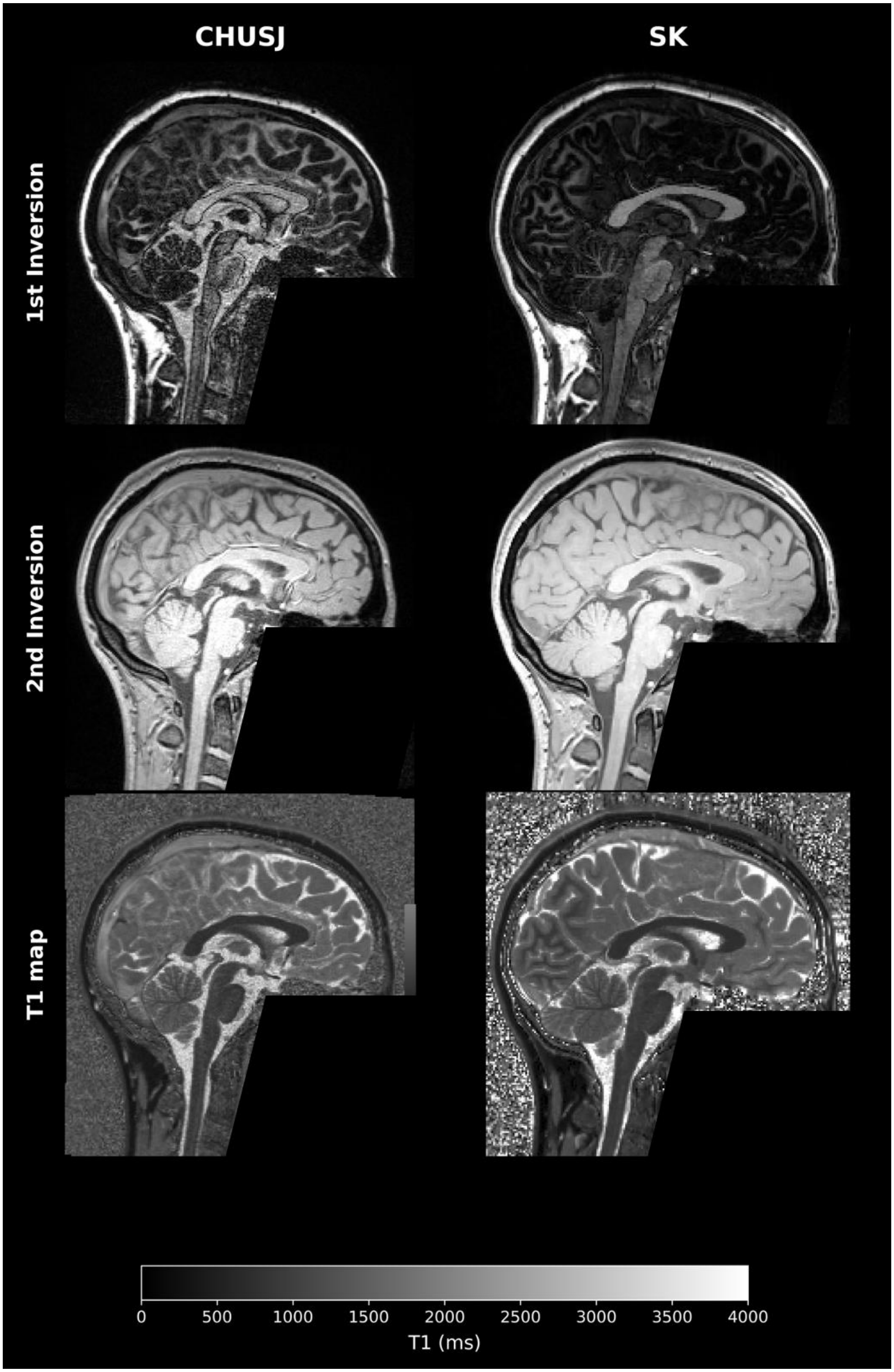
CHUSJ
SK
1st Inversion
2nd Inversion
T1 map
0
500
1000
1500
2000
2500
3000
3500
4000
T1 (ms)

**Supplementary Figure 6.** Representative sagittal slices of Magnetization prepared 2-rapid gradient echo imaging (MP2RAGE) data acquired from a single adult travelling participant (age 31) at each of the two C-PIP sites: (Left) Centre hospitalier universitaire Sainte-Justine (CHUSJ); (Right) Hospital for Sick Children – SickKids (SK). Only the quantitative T1 map is shown. Top row: Raw 1$^{st}$ inversion image; Middle row: Raw 2$^{nd}$ inversion image; Bottom row: Quantitative T1 map. Data have been defaced to protect the identity of the participant.

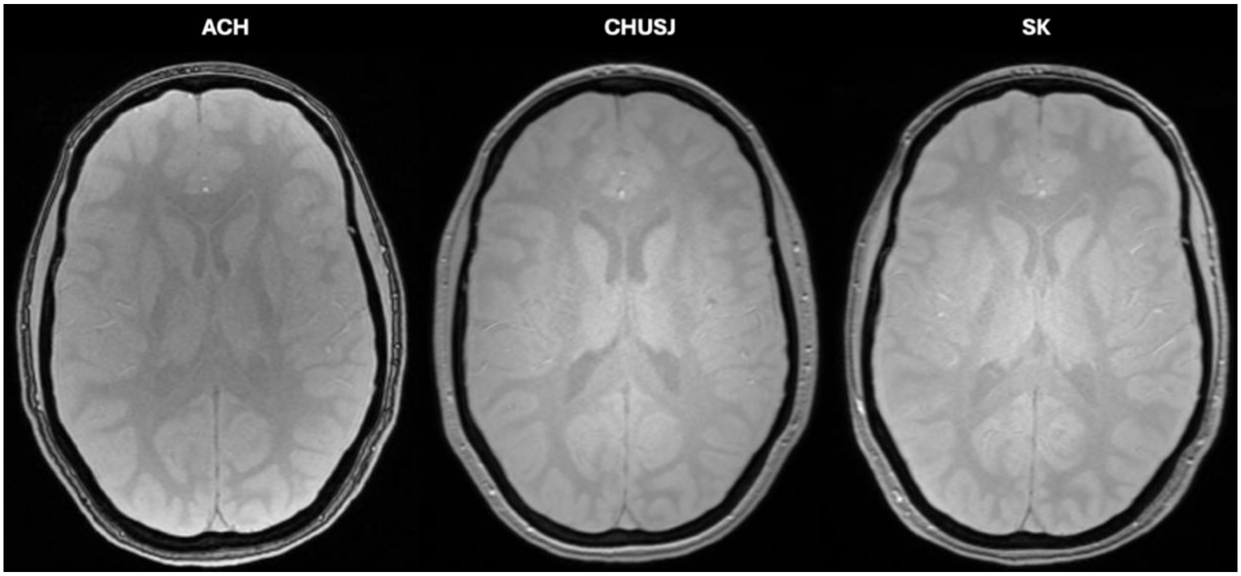


**Supplementary Figure 7.** Representative axial slices of Strategically Acquired Gradient Echo (STAGE) imaging data acquired from a single adult travelling participant (age 31) at each of the three C-PIP sites: (Left panel) Alberta Children's Hospital (ACH); (Middle Panel) Centre hospitalier universitaire Sainte-Justine (CHUSJ); (Right Panel) Hospital for Sick Children – SickKids (SK). Only the first echo of the 5° flip-angle run is shown.

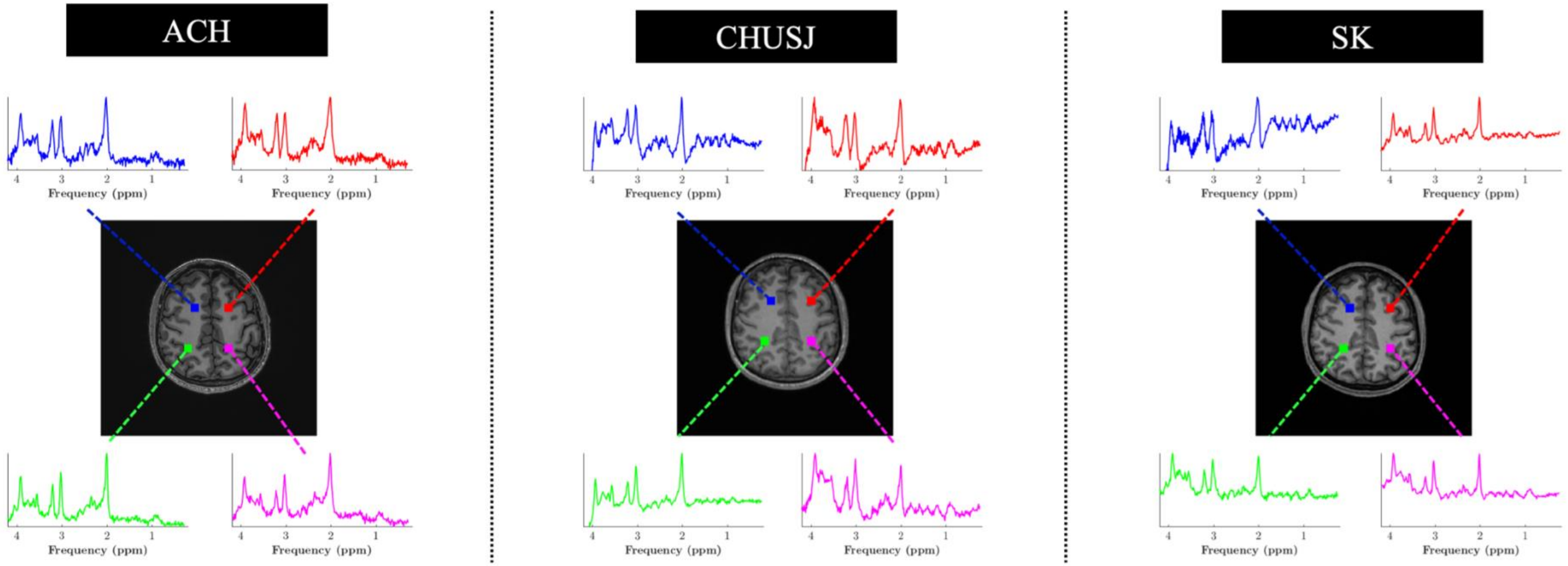


**Supplementary Figure 8.** Representative images of axial slice magnetic resonance spectroscopic imaging (MRSI) data showing the spatial location of 4 example voxels and their associated spectra acquired from a single adult travelling participant (age 34) at each of the three C-PIP sites: (Left) Alberta Children's Hospital (ACH); (Middle) Centre hospitalier universitaire Sainte-Justine (CHUSJ); (Right) Hospital for Sick Children – SickKids (SK).

**Table S1.** Censoring of high motion (>0.2 mm) ME-fMRI volumes.

| subject | site | session | movie | volumes | removed | % removed |
|---|---|---|---|---|---|---|
| Subject 1 | ACH | ses-1a | partlycloudy | 137 | 0 | 0.0 |
| Subject 1 | ACH | ses-1a | laluna | 181 | 9 | 5.0 |
| Subject 1 | ACH | ses-1b | laluna | 181 | 2 | 1.1 |
| Subject 1 | ACH | ses-1b | partlycloudy | 137 | 0 | 0.0 |
| Subject 1 | CHUSJ | ses-1a | partlycloudy | 137 | 0 | 0.0 |
| Subject 1 | CHUSJ | ses-1a | laluna | 181 | 0 | 0.0 |
| Subject 1 | CHUSJ | ses-1b | partlycloudy | 137 | 0 | 0.0 |
| Subject 1 | CHUSJ | ses-1b | laluna | 181 | 4 | 2.2 |
| Subject 1 | SK | ses-1a | laluna | 181 | 2 | 1.1 |
| Subject 1 | SK | ses-1a | partlycloudy | 137 | 0 | 0.0 |
| Subject 1 | SK | ses-1b | partlycloudy | 137 | 0 | 0.0 |
| Subject 1 | SK | ses-1b | laluna | 181 | 3 | 1.7 |
| Subject 2 | ACH | ses-1a | laluna | 181 | 1 | 0.6 |
| Subject 2 | ACH | ses-1a | partlycloudy | 137 | 3 | 2.2 |
| Subject 2 | ACH | ses-1b | partlycloudy | 137 | 2 | 1.5 |
| Subject 2 | ACH | ses-1b | laluna | 181 | 4 | 2.2 |
| Subject 2 | CHUSJ | ses-1a | laluna | 181 | 0 | 0.0 |
| Subject 2 | CHUSJ | ses-1a | partlycloudy | 137 | 0 | 0.0 |
| Subject 2 | CHUSJ | ses-1b | laluna | 181 | 16 | 8.8 |
| Subject 2 | CHUSJ | ses-1b | partlycloudy | 137 | 9 | 6.6 |
| Subject 2 | SK | ses-1a | partlycloudy | 137 | 0 | 0.0 |
| Subject 2 | SK | ses-1a | laluna | 181 | 17 | 9.4 |
| Subject 2 | SK | ses-1b | laluna | 181 | 1 | 0.6 |
| Subject 2 | SK | ses-1b | partlycloudy | 137 | 0 | 0.0 |
| Subject 3 | ACH | ses-1a | laluna | 181 | 0 | 0.0 |
| Subject 3 | ACH | ses-1a | partlycloudy | 137 | 0 | 0.0 |
| Subject 3 | ACH | ses-1b | partlycloudy | 137 | 1 | 0.7 |
| Subject 3 | ACH | ses-1b | laluna | 181 | 3 | 1.7 |
| Subject 3 | CHUSJ | ses-1a | laluna | 181 | 0 | 0.0 |
| Subject 3 | CHUSJ | ses-1a | partlycloudy | 137 | 0 | 0.0 |
| Subject 3 | CHUSJ | ses-1b | partlycloudy | 137 | 0 | 0.0 |
| Subject 3 | CHUSJ | ses-1b | laluna | 181 | 0 | 0.0 |
| Subject 3 | SK | ses-1a | partlycloudy | 137 | 0 | 0.0 |
| Subject 3 | SK | ses-1a | laluna | 181 | 0 | 0.0 |
| Subject 3 | SK | ses-1b | partlycloudy | 137 | 0 | 0.0 |
| Subject 3 | SK | ses-1b | laluna | 181 | 0 | 0.0 |

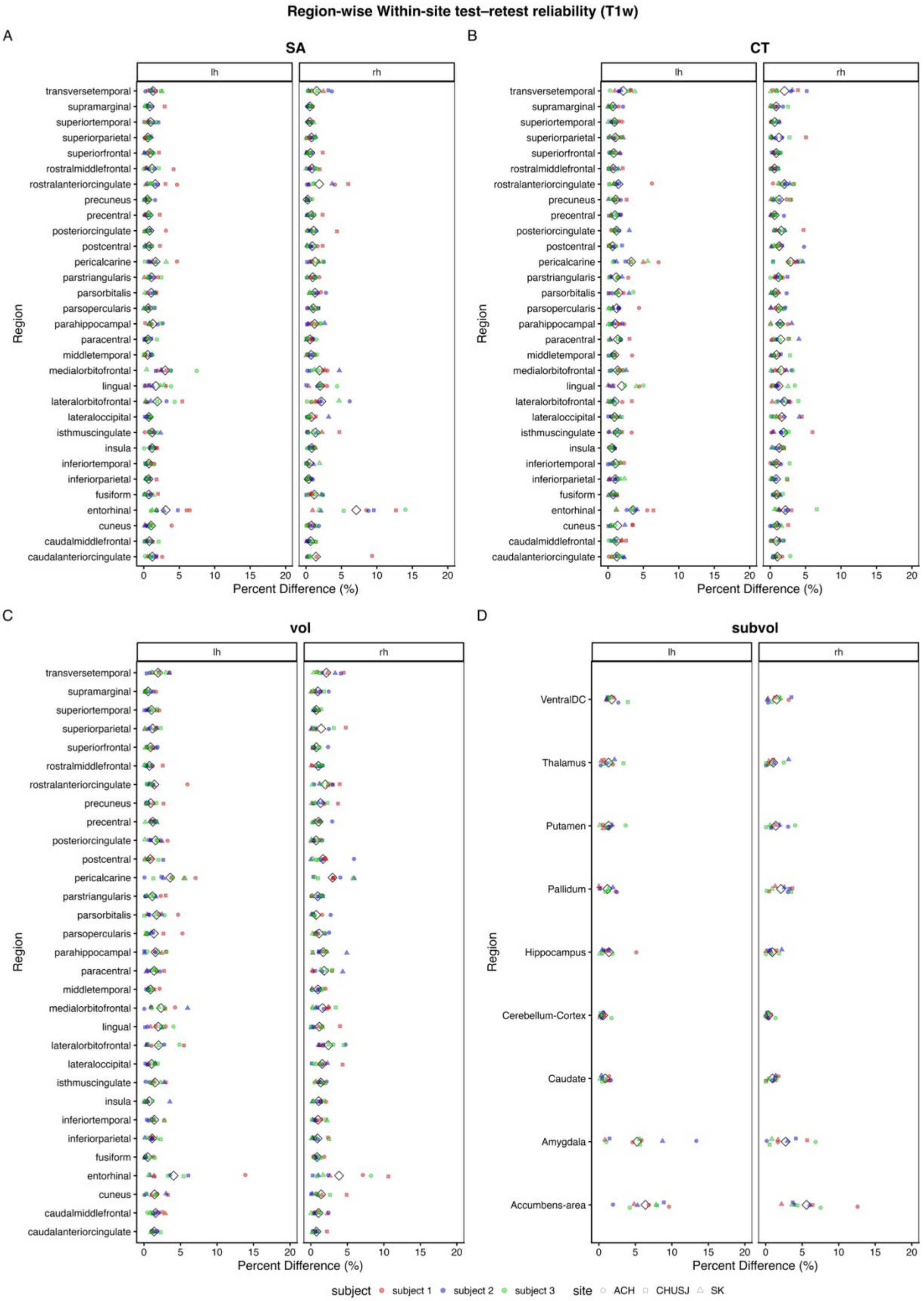

Region-wise Within-site test–retest reliability (T1w)
A
SA
B
CT
C
vol
D
subvol
lh
rh
Region
Percent Difference (%)
transversetemporal
supramarginal
superiortemporal
superiorparietal
superiorfrontal
rostralmiddlefrontal
rostralanteriorcingulate
precuneus
precentral
posteriorcingulate
postcentral
pericalcarine
parstriangularis
parsorbitalis
parsopercularis
parahippocampal
paracentral
middletemporal
medialorbitofrontal
lingual
lateralorbitofrontal
lateraloccipital
isthmuscingulate
insula
inferiortemporal
inferiorparietal
fusiform
entorhinal
cuneus
caudalmiddlefrontal
caudalanteriorcingulate
VentralDC
Thalamus
Putamen
Pallidum
Hippocampus
Cerebellum-Cortex
Caudate
Amygdala
Accumbens-area
0 5 10 15 20
subject
subject 1
subject 2
subject 3
site
ACH
CHUSJ
SK

**Supplementary Figure 9.** Absolute percentage differences between repeated sessions within each region of interest for **(A)** Surface area (SA), **(B)** Cortical thickness (CT), **(C)** Cortical volume (vol), **(D)** Subcortical volume (subvol) for left (lh) and right (rh) hemispheres. **ACH** (Alberta Children's Hospital); **CHUSJ** (Centre hospitalier universitaire Sainte-Justine); **SK** (Hospital for Sick Children – SickKids). **Subject-1**: Red; **Subject-2**: Blue; **Subject-3**: Green. **ACH**: Circle; **CHUSJ**: Square; **SK**: Triangle. White diamonds denote the average value for that region across subjects.

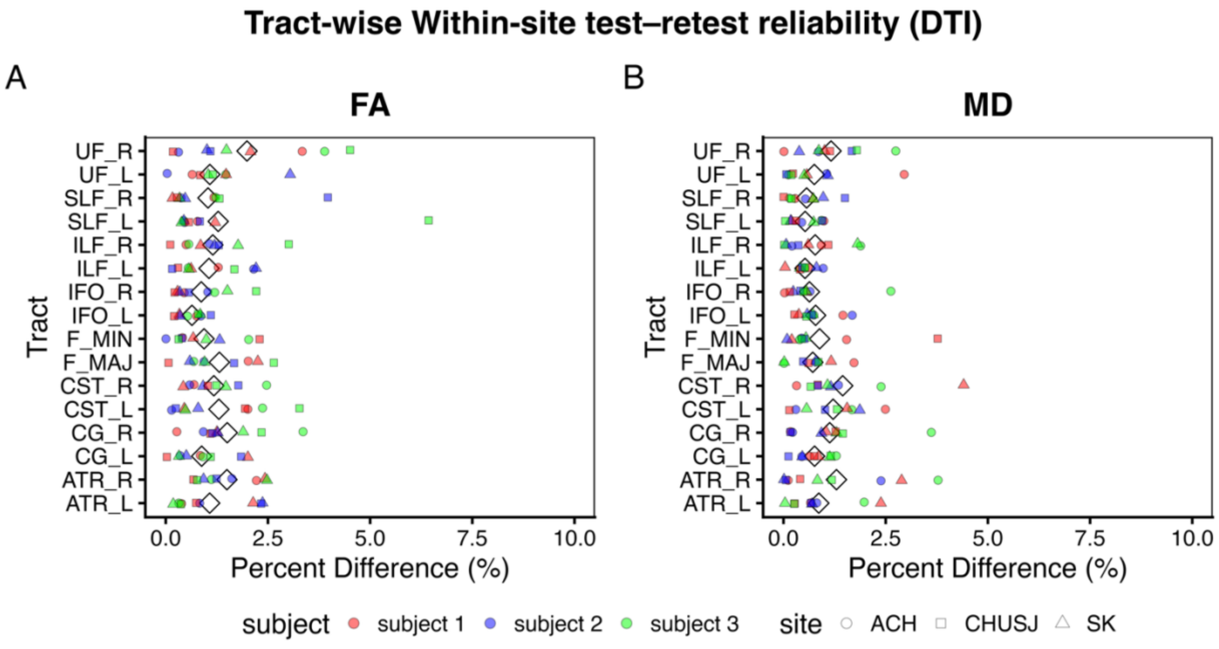


**Supplementary Figure 10.** Absolute percentage differences between repeated sessions within each tract of interest for **(A)** fractional anisotropy (FA) and **(B)** mean diffusivity (MD). **ACH** (Alberta Children's Hospital); **CHUSJ** (Centre hospitalier universitaire Sainte-Justine); **SK** (Hospital for Sick Children – SickKids). **ATR** (Anterior thalamic radiation); **CST** (Corticospinal tracts); **F_MAJ & F_MIN** (Forceps major and minor); **CG** (Cingulum); **IFO** (Inferior fronto-occipital fasciculus); **ILF** (Inferior longitudinal fasciculus); **SLF** (Superior longitudinal fasciculus); **UF** (Uncinate fasciculus). **Subject-1**: Red; **Subject-2**: Blue; **Subject-3**: Green. **ACH**: Circle; **CHUSJ**: Square; **SK**: Triangle. White diamonds denote the average value for that tract across subjects.
